# Chemogenetic Stimulation of Tonic Locus Coeruleus Activity Strengthens the Default Mode Network

**DOI:** 10.1101/2021.10.28.463794

**Authors:** Esteban A. Oyarzabal, Li-Ming Hsu, Manasmita Das, Tzu-Hao Harry Chao, Jingheng Zhou, Sheng Song, Weiting Zhang, Kathleen G. Smith, Natale R. Sciolino, Irina Y. Evsyukova, Hong Yuan, Sung-Ho Lee, Guohong Cui, Patricia Jensen, Yen-Yu Ian Shih

## Abstract

The default mode network (DMN) of the brain is involved in cognition, emotion regulation, impulsivity, and balancing between internally and externally focused states. DMN dysregulation has been implicated in several neurological and neuropsychiatric disorders. In this study, we used functional magnetic resonance imaging (fMRI), positron emission tomography (PET), and spectral fiber-photometry to investigate the selective neuromodulatory effect of norepinephrine (NE)-releasing noradrenergic neurons in the locus coeruleus (LC) on the DMN in mice. Chemogenetic-induced tonic LC-NE activity decreased cerebral blood volume (CBV) and glucose uptake, and increased synchronous low frequency fMRI activity within the frontal cortices of the DMN. Fiber-photometry results corroborated these findings, showing that LC-NE activation induced NE release, enhanced calcium-weighted neuronal spiking, and reduced CBV in the anterior cingulate cortex. These data suggest that LC-NE alters conventional stimulus-evoked coupling between neuronal activity and CBV in the frontal DMN. We also demonstrated that chemogenetic activation of LC-NE neurons strengthened functional connectivity within the frontal DMN, and this effect was causally mediated by reduced modulatory inputs from retrosplenial and hippocampal regions to the association cortices of the DMN.

## Introduction

Functional magnetic resonance imaging (fMRI) has been widely utilized to demonstrate the presence of spatiotemporally consistent intrinsic functional brain networks during resting-state. The default mode network (DMN), comprised of the prefrontal, orbitofrontal, prelimbic, cingulate, retrosplenial, posterior parietal, and temporal association cortices as well as the dorsal hippocampus, is among the most robust intrinsic networks due to its highly synchronized activity in the absence of cognitive tasks or saliency (*1*). The DMN is vulnerable in several neurological and neuropsychiatric disorders (*2*), is functionally associated with a wide range of behaviors (*3*), integrates interoceptive and exteroceptive information from multiple brain networks (*4*) and maintains the brain in a semi-vigilant state (*5*). In order to make causal interpretations of behaviorally relevant DMN changes and design network-based interventions for disorders that afflict DMN activity, identifying the modulatory mechanisms controlling the DMN is of paramount importance.

The locus coeruleus (LC), a small nucleus within the pons, is a potential DMN modulator (*6, 7*). A large portion of the neuromodulator norepinephrine (NE) originates from the LC and is released in the brain regions that are considered DMN nodes (*8, 9*). Accumulating evidence suggests that LC-NE may be essential for DMN modulation since: **a)** NE receptors are prominently expressed in DMN-related brain structures (*9*); **b)** LC-NE can bidirectionally modulate attention reorientation in a dose-dependent manner (*8*); **c)** LC-NE neuron degeneration and DMN disruption are coincidently found in depression (*10*), traumatic brain injury (*11*), Parkinson’s disease (*12*), Alzheimer’s disease (*13*) and aging (*14*); and **d)** pharmacological treatment of pathological LC-NE levels reduces attentional lapses (*15*) and restores DMN integrity in attention deficit hyperactivity disorder patients (*16*). Despite these findings, the modulatory association between LC-NE and DMN remains circumstantial since pharmacological interventions using NE-related agents inherently result in non-selective binding on dopaminergic, cholinergic and serotonergic receptors (*7, 17, 18*). Furthermore, systemic administration of these agents indiscriminately targets all NE-producing neurons in the brain and sympathetic nervous system, making it difficult determine the role of the LC-NE in modulating the DMN. Though selective manipulation of LC-NE while imaging DMN is currently impossible in humans due to technical and ethical constraints (*7, 19*), such studies are feasible in rodent models since structural and functional homologs of the human DMN have been identified in mice (*20*–*27*).

In this study, we used an established data-driven approach to identify DMN modules (*28*) and an intersectional chemogenetic strategy to selectively and reproducibly induce tonic LC-NE activity in mice (*29, 30*) to causally determine a circuit model of this modulation. To reveal potential confounders that could impact our interpretations of LC-NE influence on the DMN, we measured changes in several fMRI metrics, neuronal calcium activity, and glucose uptake across different spatial and temporal scales. Through modeling the signal dynamics, we revealed the circuit mechanism by which LC-NE activation modulates the DMN. Our findings should pave the way towards a better understanding of how large-scale brain networks are mediated by a specific neuromodulatory system.

## RESULTS

To selectively and reproducibly activate LC-NE neurons, we used an intersectional chemogenetic approach in which the excitatory G-protein-coupled receptor hM3Dq, fused to mCherry, is expressed in 99.6% of the anatomically defined LC located within the central gray and a small portion of the dorsal subcoeruleus immediately adjacent to and continuous with the LC (LC-NE/hM3Dq) (*30*) (**Fig. 1**). This population of NE neurons robustly innervates canonical DMN regions including cingulate 1 (Cg1) and retrosplenial (RSC) cortices (**fig. S1**) (*31*).

**Fig 1.**
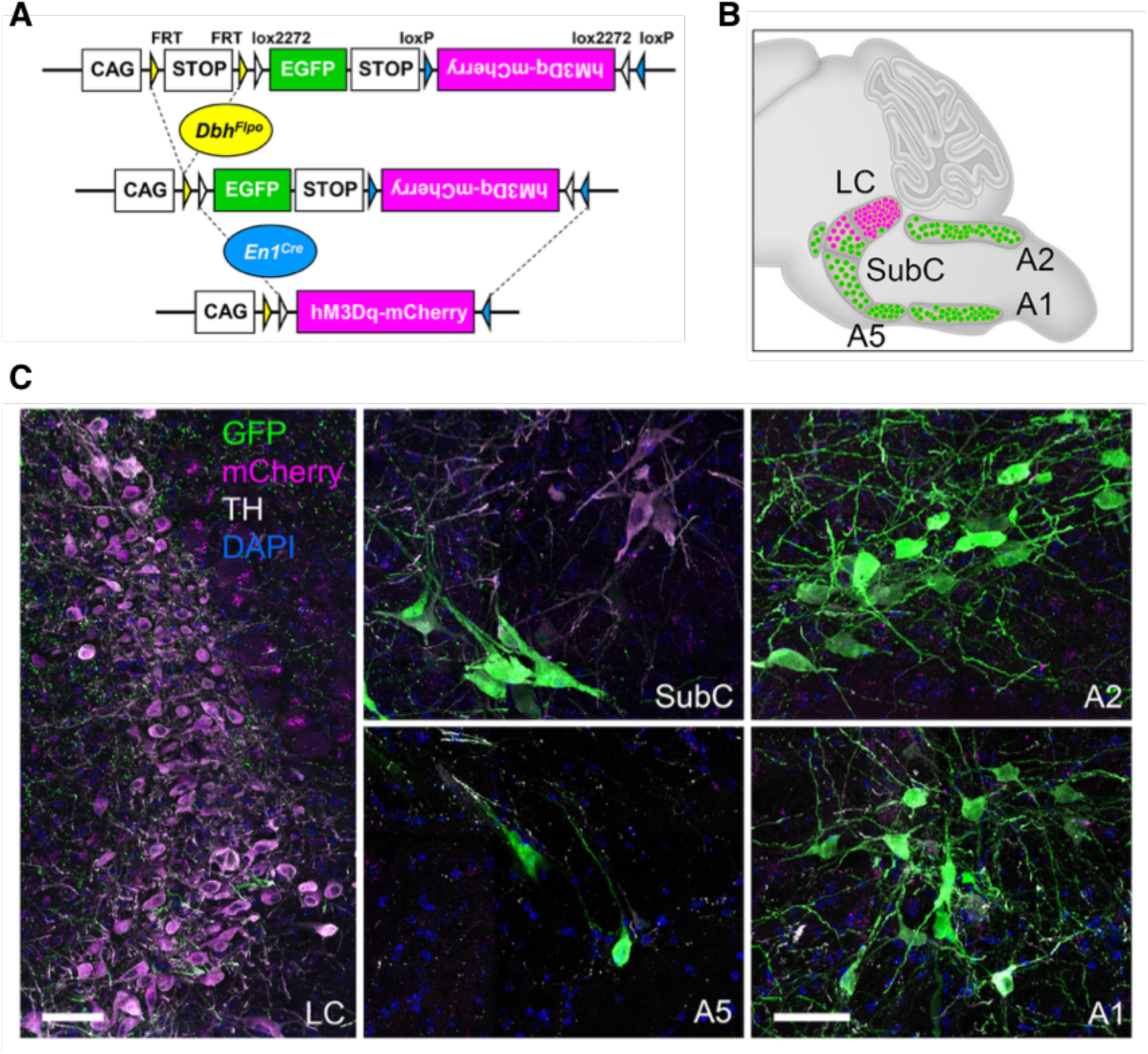
Intersectional chemogenetic strategy to selectively activate LC-NE neurons. **(A)** Schematic illustration of the intersectional genetic strategy. **(B)** A sagittal schematic diagram of the hindbrain compressed along the mediolateral axis illustrates the approximate position of NE neurons. Recombination of the RC::FL-hM3Dq allele by the noradrenergic-specific driver *Dbh*^*Flpo*^ and *En1*^*cre*^ results in expression of the excitatory G-protein-coupled receptor hM3Dq fused to mCherry in LC-NE neurons (magenta cells in schematic). Expression of *Dbh*^*Flp*^ by all remaining NE neurons results in expression of GFP (green neurons in schematic). **(C)** Immunofluorescent labeling of sections from the adult brainstem of LC-NE/hM3Dq mice reveals hM3Dq-mCherry-expressing NE neurons in the LC (magenta) and GFP-expressing NE neurons (green) in the SubC, A5, and C2/A2, and C1/A1 nuclei. Scale bar indicates 50 µm.

To functionally delineate the DMN and study selective NE modulatory effects, we performed *in vivo* CBV-weighted fMRI scans on LC-NE/hM3Dq mice and littermate controls (n=9 and 12, respectively) under light isoflurane (∼1%) anesthesia using a previously described isotropic echo-planar-imaging (EPI) protocol (*32*). Though medetomidine with low dose isoflurane is considered the preferred sedative for rodent fMRI (*33*), we avoided its usage since it suppresses NE release (*34*). We collected a 10 min resting-state baseline scan prior to administering a 1 mg/kg intraperitoneal injection of clozapine-n-oxide (CNO) (**Fig. 2A**). This protocol has been shown to activate LC-NE neurons at tonic frequency and suppress locomotion in LC-NE/hM3Dq mice (*29, 30*). Subsequent comparisons were made against littermate controls to account for off-target effects of CNO and/or the back-metabolized clozapine (*35*). We spatially warped each imaging dataset into the Allen Mouse Common Coordinate Framework (**fig. S2**), functionally parcellated the baseline fMRI data from all subjects (*n*=21) by performing a 100-component independent component analysis (ICA) (**Fig. 2, B and C**), and verified their reproducibility (**fig. S3**). We identified 17 DMN-related independent components (ICs) (**fig. S4A**) according to previous rodent DMN studies (*22, 24, 25, 28, 36*). The areas showing significant temporal correlation associated with the 17 identified DMN ICs were reconstructed using dual regression, and a one-sample two-sided *t*-test was performed to generate the group-level maps representing the connectivity of these ICs. These maps showed high spatial similarity with an RSC seed-based connectivity map commonly used to depict the DMN (**fig. S4, B and C**). Louvain community modularity analysis clustered the 17 DMN ICs into three distinct modules (Q = 0.10, p<0.01; **Fig. 2D**): A *Frontal* module comprised of pre-limbic/infra-limbic (PrL/IL), lateral orbital (LO), Cg1, and anterior cingulate 2 (aCg2) cortices; an *RSC-HIPP* module comprised of the dorsal hippocampus (HIPP) and posterior Cg2 (pCg2), medial parietal association (MPtA), retrosplenial granular (RSG), retrosplenial dysgranual / visual (RSD / Vis) cortices; and an *Association* module comprised of posterior parietal (PPtC), and auditory (Aud) cortices. No significant difference among the connectivity of DMN ICs were found in the pre-CNO baseline data between LC-NE/hM3Dq and control groups (*P*_*FDR-corrected*_ > 0.05, **fig. S4D**).

**Fig. 2.**
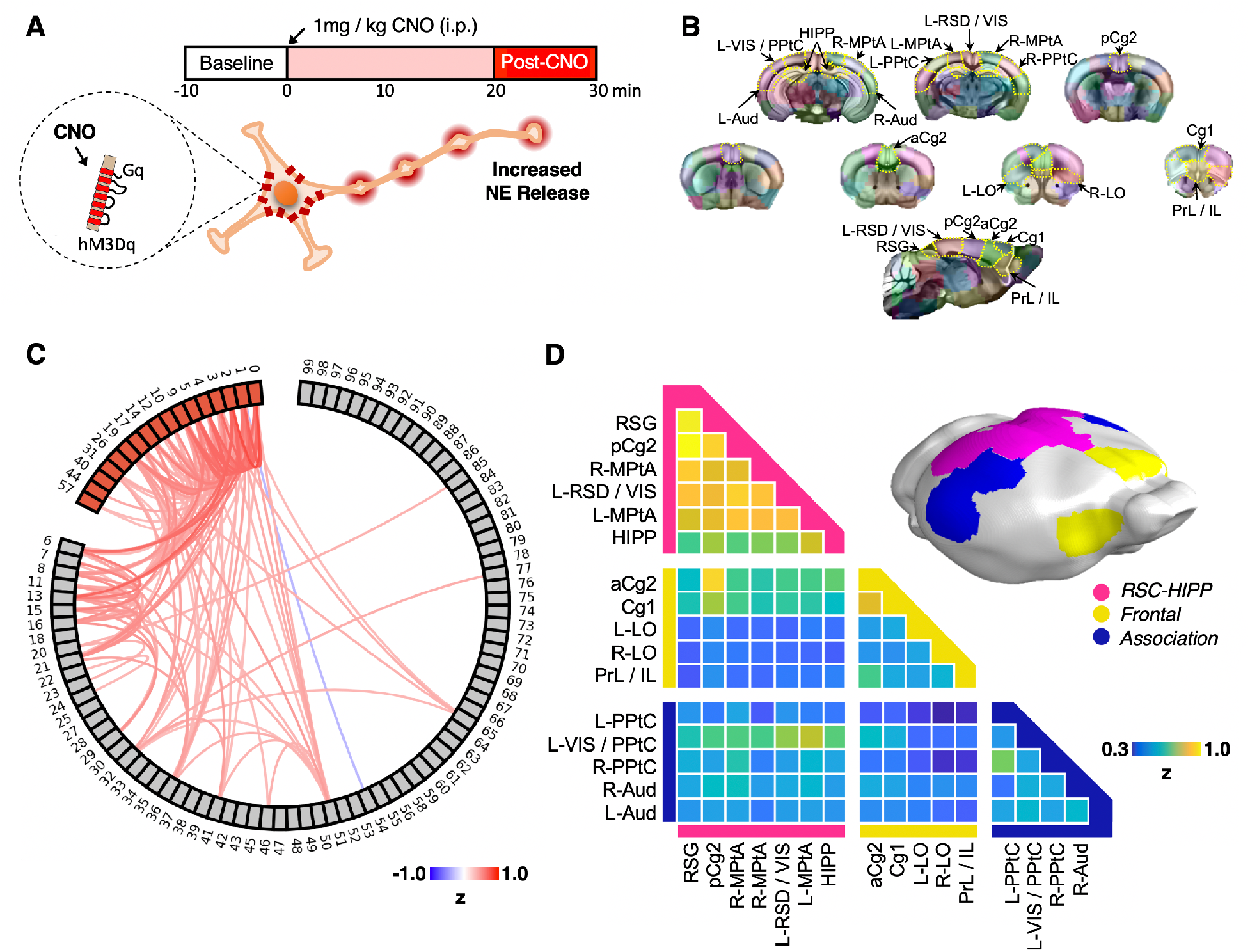
Experimental design and identification of mouse DMN modules. **(A)** CBV fMRI experimental design includes a 10 min baseline scan prior to CNO administration and 30 min of scans after CNO. **(B)** 100 ICs were derived from group ICA of baseline scans among all subjects. Specific IC masks were determined by a winner-take-all strategy through comparing mean z-values from all ICs on a voxel basis and then color-coded. DMN-related ICs were then identified according to rodent DMN topology in the literature. **(C)** Correlation among the 100 ICs were plotted with a threshold Fischer | z | > 0.3. DMN ICs were labeled in red. **(D)** Modularity analysis of DMN ICs showing that the mouse DMN is comprised of Frontal (yellow), *RSC-HIPP* (pink), and *Association* modules (blue).

Using the *Frontal, RSC-HIPP, and Association* module ROIs (one-sample two-sided *t*-test, *P* < 0.0001) and striatum as a reference region due to sparse innervation from LC-NE neurons, we examined how NE release from LC modulates fMRI-derived CBV, regional homogeneity (ReHo) and amplitude of low frequency fluctuation (ALFF) changes. We compared the 10 min, pre-CNO baseline fMRI data against data acquired between 20-30 min post-CNO to account for CNO-induced activity kinetics reported in fMRI (*6, 37*–*41*) and behavior studies (*29, 30*). CNO-evoked LC-NE activation significantly decreased CBV in all DMN modules in LC-NE/hM3Dq (unpaired *t* test, *P*_*FDR-corrected*_ < 0.05) but not in control mice (**Fig. 3A**). Interestingly, LC-NE activation induced robust CBV changes in the striatum despite sparse innervation from LC-NE neurons (*6*), possibly due to NE-induced vasoconstriction at watershed arteries upstream of the striatum (*42*). While CBV changes appeared less specific, ReHo (**Fig. 3B**) and ALFF (**Fig. 3C**) changes were more localized, and the increase of these signals contradicted the intuitive interpretation of CBV by implying a possible increase of synchronous, low-frequency activity in the *Frontal* and *RSC-HIPP* DMN modules by LC-NE. Specifically, LC-NE activation enhanced ALFF changes within 0.01 and 0.05 Hz band frequencies (unpaired *t* test, *P*_*FDR-corrected*_ < 0.05; **fig. S5A**).

**Fig. 3.**
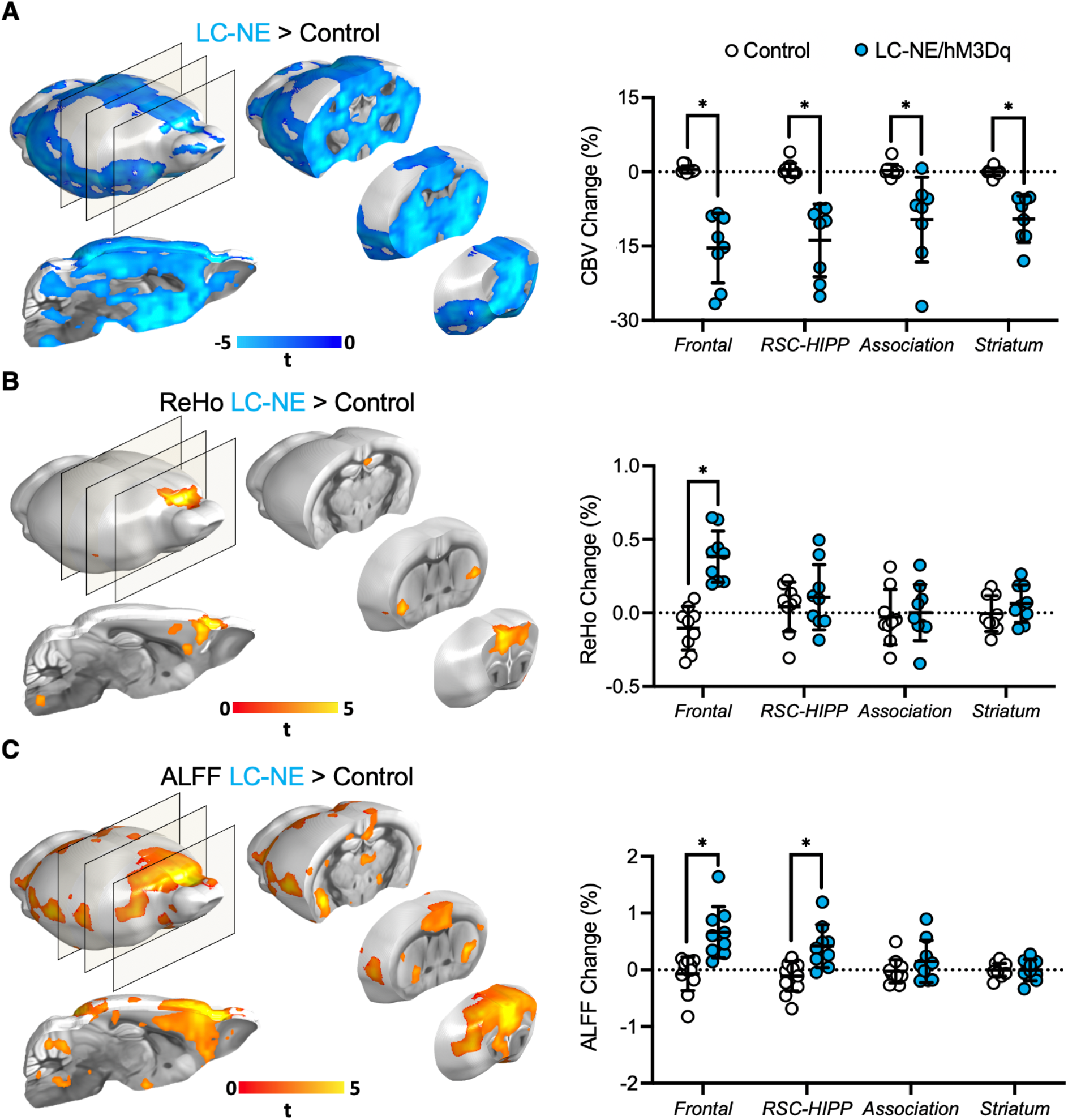
CBV, ReHo, and ALFF changes in DMN modules following activation of LC-NE neurons. **(A)** CBV decreased significantly following LC-NE release across all DMN modules compared to baseline and controls. LC-NE release significantly increased **(B)** ReHo in the *Frontal* DMN module and **(C)** ALFF in *Frontal* and *RSC-HIPP* modules compared to baseline and controls. * p < 0.05, ** p < 0.0001, horizontal lines represent means, and error bars represent ± SD.

To validate these fMRI findings, we employed spectral fiber-photometry (**Fig. 4A**) in LC-NE/hM3Dq (n=5) and control mice (n=4). We virally expressed a genetically engineered NE_2.1_ sensor (*43*) and a red-shifted jRGECO1a calcium activity sensor (*44*) under the pan-neuronal human Synapsin-1 (hSyn) promoter in the Cg1 of the *Frontal* DMN module (**Fig. 4B**), where the effects of LC-NE activation were most robust, and intravenously administered a CY5-conjugated Dextran far-red fluorescent dye to measure CBV (**fig. S6**). Collectively, this allowed us to simultaneously detect changes in synaptic NE release, neuronal activity-mediated calcium influx, and CBV in Cg1 (**Fig. 4C**). After CNO administration, we observed significant increases in NE release (unpaired t test, *P*_FDR-corrected_ < 0.05; **Fig. 4D**), decreases in CBV (unpaired t test, *P*_FDR-corrected_ < 0.05; **Fig. 4E**), and increases in the neuronal activity (unpaired t test, *P*_FDR-corrected_ < 0.05; **Fig. 4F**) of LC-NE/hM3Dq mice compared to controls. Notably, the number of calcium spikes also increased significantly following CNO administration (unpaired t test, *P*_FDR-corrected_ < 0.05; **Fig. 4G**), a hallmark of NE that tunes the signal-to-noise ratio of downstream neuronal firing (*45*). These findings corroborate well with our fMRI results (**Fig. 3**) and indicate that driving LC-NE release can concurrently induce regional vasoconstriction while enhancing neuronal excitability in Cg1. Given that fMRI does not directly measure neuronal activity, these results highlight the importance to cautiously interpret fMRI-derived DMN results when NE is involved, as inferring neuronal activity by direct fMRI signal changes in this case may be erroneous.

**Fig. 4.**
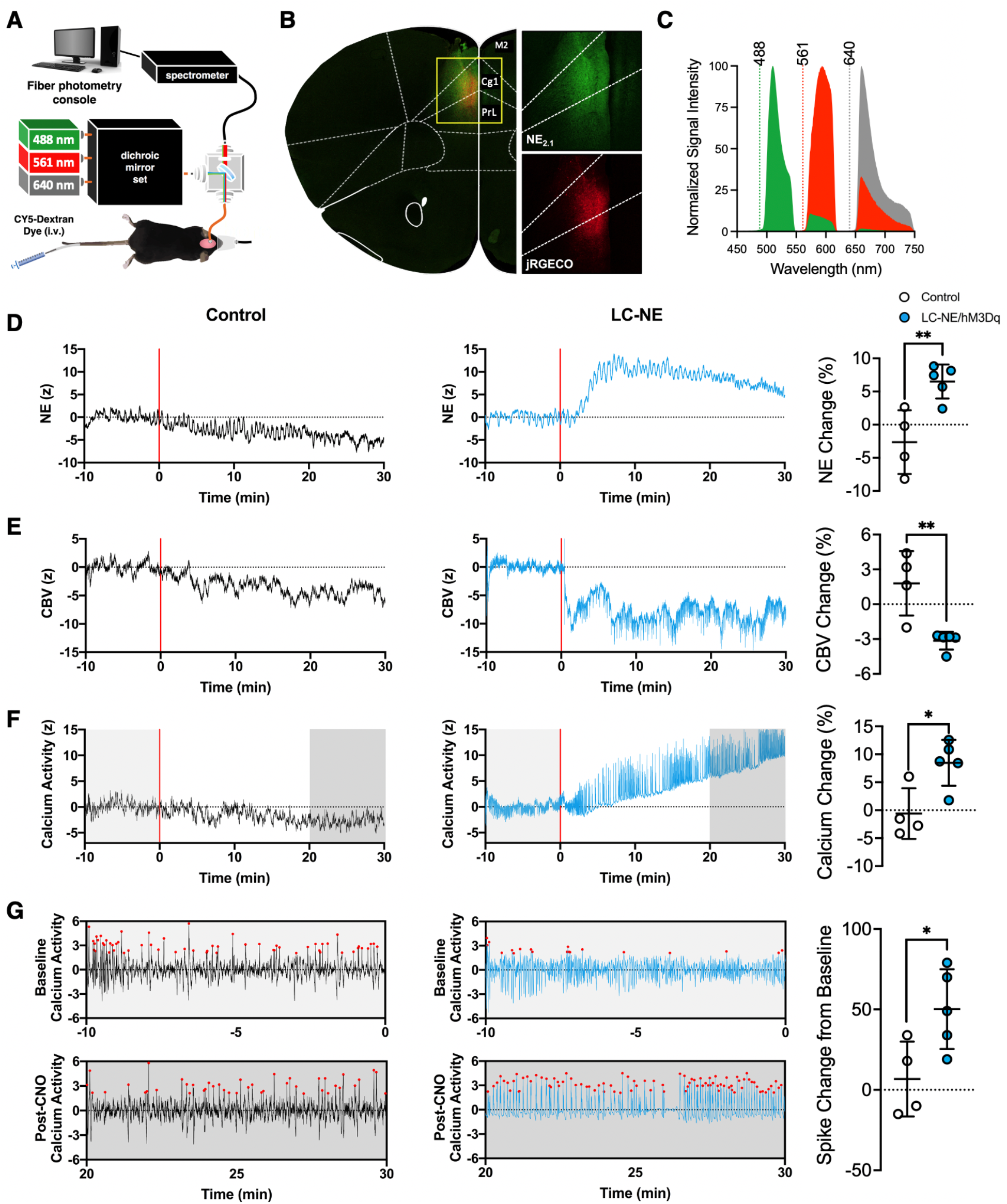
Triple-spectral fiber photometry measuring NE release, CBV, and calcium-weighted neuronal activity in Cg1. **(A)** Experimental setup of the fiber-photometry system with 488, 561 and 640 nm lasers simultaneously used to detect NE release (NE_2.1_), neuronal calcium activity (jRGECO1a), and CBV (CY5-Dextran dye) changes, respectively. **(B)** Cg1 neurons were confirmed to be transfected to express NE_2.1_ (green) and jRGECO1a (red) sensors. **(C)** Spectral profiles of NE neuronal activity and CBV sensor emissions used to resolve signals via an established spectral unmined approach. **(D-F)** Respective effects of CNO on NE release, CBV, and neuronal activity in Cg1 from representative subjects and group level bar graphs. Subjects were continuously scanned for 40 min with a 1 mg/kg dose of CNO administered via an intraperitoneal catheter 10 min (t = 0) after scan onset. **(G)** CNO-induced LC-NE activation altered post-synaptic calcium spiking patterns, resulting in a significant increase in spike counts. Red dots represent the Ca^2+^ spikes (z-score > 1.96). * *p* < 0.05, ** *p* < 0.01, horizontal lines represent means, and error bars represent ± SD.

We also examined the effect of NE on brain glucose uptake in a subset of LC-NE/hM3Dq (n=5) and control (n=5) mice using an established [^18^F]-fluorodeoxyglucose (FDG)-PET protocol (*46*). Interestingly, we found that CNO-induced LC-NE activation significantly decreased glucose uptake in all three DMN modules (paired *t* test, *P*_*FDR-corrected*_ < 0.001), but not in control subjects (**Fig. 5** and **S7**). Though there are mixed findings on how NE alters glucose uptake (*47*–*52*), most studies using pharmacology to enhance NE release showed suppressed glucose uptake (*47*–*52*) by shifting astrocyte metabolism to preferentially utilize glycogen reserves (*50, 51*). Taken together, these findings suggest additional caution needs to be considered for FDG-PET data interpretation when triggering NE release.

**Fig. 5.**
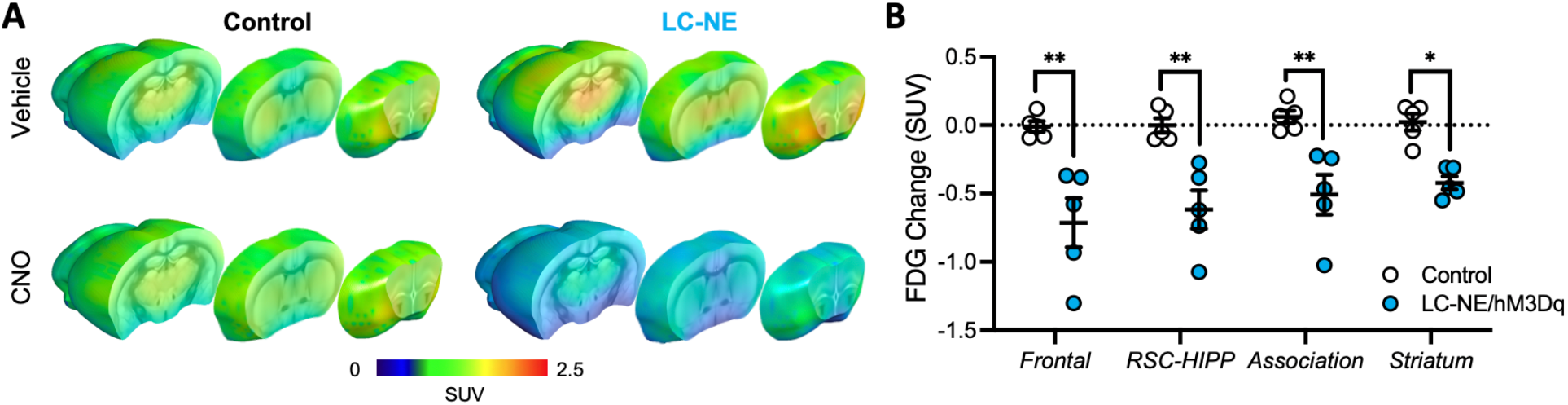
Glucose uptake changes in DMN modules brain wide following activation of LC-NE neurons. **(A)** Activation of LC-NE neurons significantly decreased FDG uptake compared to saline-treated sham and littermate controls receiving CNO. **(B)** LC-NE activation significantly reduced standard uptake values (SUV) in all DMN regions. SUV changes were derived from each subject underwent two scans (vehicle and CNO administration) following co-registration. Comparisons were made against littermate controls. * *p* < 0.001, ** *p* < 0.0001, horizontal lines present means, and error bars represent ± SD.

To access how selective NE release modulates DMN connectivity and network properties, we examined FC changes within and between *Frontal, RSC-HIPP*, and *Association* DMN modules in LC-NE/hM3Dq (n=9) and control mice (n=12). We found that CNO significantly enhanced FC within *Frontal* (F_(2,26)_=5.2, *P*_*FDR-corrected*_ < 0.05) and between *Frontal* and *Association* modules (F_(2,26)_ =6.0, *P*_*FDR-corrected*_ < 0.05) of the DMN in LC-NE/hM3Dq, but not in control mice (**Fig. 6, A and B**). Notably, the high intramodular connectivity characterized by within-module degree (WD), which quantifies the level of connectivity of nodes within a module, indicated that Cg1 and RSC serve as provincial hubs for the *Frontal* and *RSC-HIPP* modules of the DMN, respectively (**fig. S8, A and B**). The high intermodular connectivity characterized by the partition coefficient (PC), which estimates the level of interactions with nodes of other modules, indicated that the aCg2 of the *Frontal* module serves as a connector hub throughout the entire DMN and may control the FC between *Frontal* and other modules (**fig. S8, A and B**). Given the putative causal control of the anterior insular cortex (AI) on the *Frontal* module of the DMN (*26, 53*), we also examined the effects of LC-NE activation on their FC changes and found that CNO-evoked activation significantly enhanced anticorrelation between AI and DMN frontal module (**fig. S8, C and D**).

**Fig. 6.**
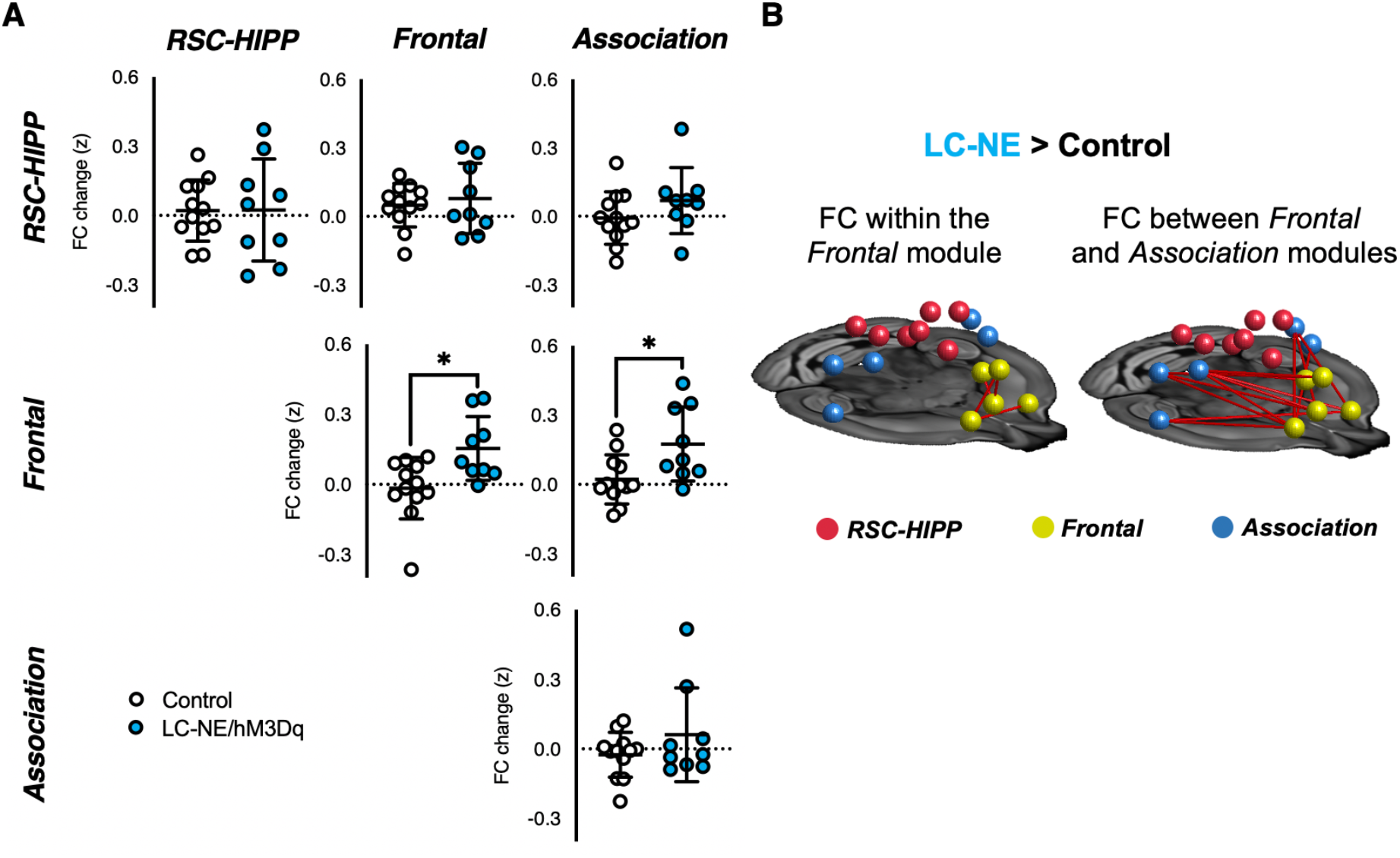
FC changes among DMN modules following activation of LC-NE neurons. **(A)** Within and between DMN module FC changes. **(B)** Network-based statistical analysis showing significant differences in FC among edges in LC-NE > Control. * *P* _FDR-corrected_ < 0.01, horizontal lines represent means, and error bars represent ± SD.

To further unravel the causal influences on FC in DMN modules by LC-NE activation, we conducted a dynamic causal modeling (DCM) analysis and found that LC-NE activation significantly reduced effective connectivity (EC) from *RSC-HIPP* to the *Association* module (paired two-sided *t*-test, *P*_*FDR-corrected*_ < 0.05, **Fig. 7A**), while no change was detected in the littermate controls (**fig. S9A**). Co-activation pattern (CAP) (*54*) analyses may support these findings, as the CAP representing distinct *RSC-HIPP* and *Association* states was also suppressed following LC-NE activation (**fig. S9B-D**). Together with the robust NE modulatory effects in the *Frontal* module, these findings prompted us to conduct a mediation analysis to determine the origin of the FC changes in the *Frontal* module. Using a general linear model (GLM), we demonstrated that the reduction in EC observed from *RSC-HIPP* to the *Association* module causally manipulates the FC changes within the *Frontal* module (coefficient = -0.53±0.13, *P* < 0.001; **Fig. 7B**). When we incorporated the FC increases between the *Frontal* and *Association* modules as a mediator, we observed a full mediation effect of the reduced EC to the *Frontal* module FC changes while the direct relationship between the reduced EC from *RSC-HIPP* to the *Association* module and the increased FC within *Frontal* module became insignificant (coefficient = -0.26±0.13, N.S.). The Sobel test further indicated the significant mediation effect (Sobel z = -2.02, *P* < 0.05; **Fig. 7B**). In contrast, we did not observe the reduced EC from the *RSC-HIPP* to *Association* module being mediated by the FC increases between *Frontal* and *Association* modules (**fig. S10**). Collectively, these findings indicate that LC-NE activation modulates the DMN by **1)** strengthening FC within the *Frontal* module, **2)** strengthening FC between *Frontal* and *Association* modules, and **3)** reducing *RSC-HIPP* control over the *Association* module, which causally alters *Frontal* module FC, with the FC between *Frontal* and *Association* modules serving as a key mediator.

**Fig. 7.**
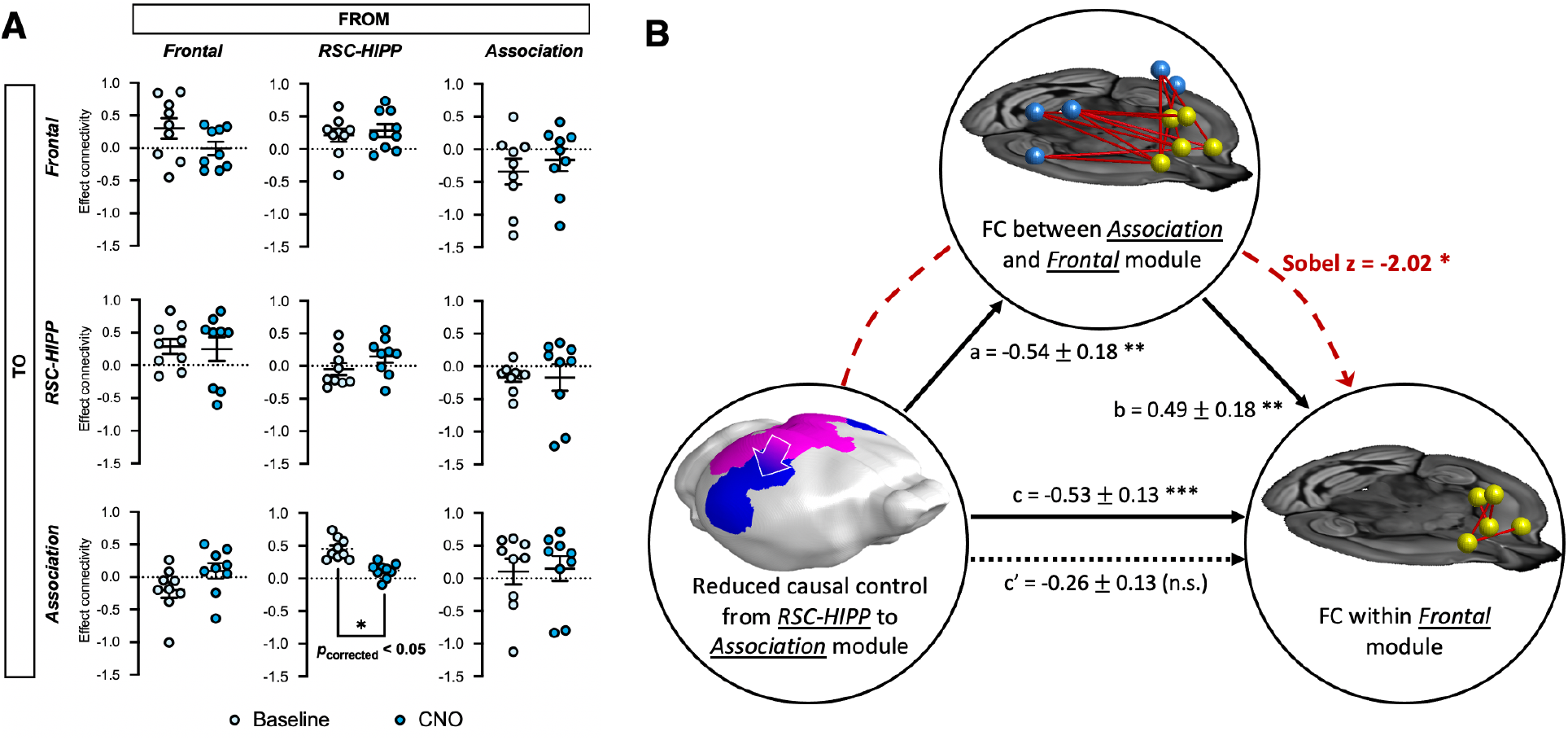
DCM and mediation analysis among DMN modules. **(A)** DCM analysis among DMN modules found that *RSC-HIPP* reduced its causal modulation to the *Association* module upon LC-NE activation (*P*_FDR-corrected_<0.05, horizontal lines represent means, and error bars represent ± SD). **(B)** Mediation analysis was performed with direct effect (a, b, c) and with a mediator (c’). A Sobel test was also performed (red dot line) to evaluate the significance of mediation effect (Sobel z-value = -2.02, *P* < 0.05). * *P* < 0.05, ** *P* < 0.01, *** *P* < 0.001, n.s. = no significance.

## DISCUSSION

The mouse DMN is well documented in the literature (*20*–*25*) and its structural foundation has been recently established by a pivotal study by Whitesell et al. (*25*). Most studies analyzing the mouse DMN using ICA or seed-based connectivity analyses report consistent DMN architectures that are homologous to the human DMN. A multi-center study compiled by Grandjean *et al*. analyzed several resting-state mouse fMRI datasets acquired under various conditions (e.g., magnetic field strengths, coils, imaging parameters, and anesthesia protocols) and generated a group ICA atlas delineating mouse brain connectivity (*22*). The results pulled three DMN modules that include prefrontal, cingulate/retrosplenial, and temporal association areas. Similar to those findings, our data analysis revealed three modules including *Frontal, RSC-HIPP*, and *Association* modules, where the *Frontal* and *RSC-HIPP* modules included prefrontal and posterior cingulate components, respectively (**Fig. 2C**). It should be noted that the involvement of the hippocampus in the mouse DMN remains disputed due to the lack of direct anatomical projections (*22, 24, 25*). This study includes hippocampus as part of the DMN because we followed established studies identifying the DMN constituents across rodents (*28, 36*) and humans (*4*). Additionally, several unbiased hierarchical clustering analyses (*24, 55*), including our own (*28, 32*), functionally classified the hippocampus as part of the DMN and lend further support to the validity of the DMN regions used for the analysis in this study.

Several research groups have pioneered chemogenetic fMRI (*6, 37*–*41*) and PET (*46*) approaches to selectively map the influence of neurotransmitters on brain networks in rodents. Our experimental design employing an intersectional chemogenetic mouse line presented a unique opportunity to investigate functional DMN modulation via a strategy that permits noninvasive and reproducible activation of LC-NE neurons (*30*). We demonstrated that chemogenetic activation of LC-NE neurons significantly reduced CBV and glucose uptake among all three DMN modules compared to littermate controls (**Fig. 3 and 5**). Utilizing littermate controls is essential for rigor in chemogenetic studies (*56*) since off-target binding of CNO or clozapine through reverse-metabolism of CNO may occur (*35*). Of note, the modulatory effect seen in the striatum suggests that LC-NE influence on these metrics may extend beyond the brain regions receiving vast innervations from LC-NE neurons.

While both fMRI and fiber-photometry corroborated the findings of CBV reduction in the *Frontal* DMN module following activation of LC-NE neurons, we observed robust increases in synchronous low frequency activity as measured by ReHo, ALFF, and photometry-derived calcium activity in our experimental condition (**Fig. 4F and G**). The most straightforward interpretation of the fMRI data based on well-documented neurovascular coupling rules (*57*) does not apply, likely due to the potent vasoconstrictive properties of LC-NE (*58*). It is not surprising that the neuromodulatory effect induced by NE release appears inconsistent in the literature owing to various brain states and basal firing rate examined, as well as the distinct experimental approaches utilized to promote/benchmark NE release (*6, 59*–*61*). The literature also shows mixed findings regarding the effects of NE on both cerebral hemodynamics and metabolism, possibly because common pharmacological agents used to induce NE release suffer from differential actions on NE receptor subtypes and non-selective binding that can either increase (*62*) or decrease perfusion (*58, 63*) and increase (*47, 52*) or decrease glucose metabolism (*48, 49*). Of note, the changes observed in glucose metabolism following LC-NE stimulation could also be attributed to dose-dependent shifting of glial metabolism, whereby high NE levels shift metabolism to preferentially glycogen deposits from blood glucose (*50, 51*). Unlike many studies that use pharmacological manipulations of NE release, our chemogenetic approach coupled with multimodal measurement of NE, neuronal activity, and CBV revealed an effect of NE in increasing synchronized low frequency activity, strengthening neuronal firing, and decreasing CBV. This has significant implications when using fMRI to interpret DMN neuronal activity, as “deactivation” of raw fMRI signal caused by LC-NE activation may not necessarily represent reduced DMN neuronal activity.

Recent animal and human fMRI studies support the role of NE in brain network reorganization (*6, 60, 61, 64*). Importantly, NE release has been shown to promote topological integration within the network (*65, 66*). This aligns well with our findings showing enhanced ReHo and FC within the *Frontal* DMN module following CNO-evoked LC-NE activation. Additionally, our ALFF results show LC-NE activation enhanced low-frequency power of the *Frontal* DMN that was restricted to a frequency band ranging from 0.01 to 0.05 Hz—a range associated with robust changes in human DMN (*67*). Alterations in the power within this sub-frequency band of ALFF also plays a vital role in attention reorientation during visual-motor attentional tasks (*67*), which has also been linked to changes in cortical NE levels (*68*).

LC-NE enhanced FC within the *Frontal* module and between *Frontal* and *Association* modules. In agreement with our finding, a seminal study virally transfecting hM3Dq via the *Dbh* promotor into the LC found significant FC enhancement in the Cg1, Cg2, and RSC following chemogenetic stimulation (*6*). Abnormally heightened FC in *Frontal* DMN regions has been associated with anxiety and depression (*69*). Anxiety-like behavioral phenotypes such as reduced locomotor activity in a novel environment and anhedonia have also been found following chemogenetic-evoked LC-NE activation in awake mice (*29*). Interestingly, such behavior response is similar to optogenetic-induced LC-NE neuronal activation at tonic frequencies (*70*). Together with our data showing tonically elevated NE release (**Fig. 4D**) and strengthened DMN connectivity (**Fig. 6**), these results suggest that tonic firing of LC-NE neurons shifts the brain towards a DMN-dominated state and therefore facilitates DMN-associated behaviors (*71*).

As we did not observe any FC decreases in the DMN and only identified strengthening FC within and between modules, our results support the functional integration theory of NE proposed by Shine (*61, 65*), complemented by the results of Zerbi et al. (*6*), and suggest that LC-NE induced functional integration could occur at a rather focal, sub-network level within the DMN. Additionally, we found FC strengthened anticorrelated coupling between the *Frontal* DMN module and AI, a key node of the salience network (SN) that may causally suppress the DMN (*53*) (**fig. S8, C and D**). This also indicates that integration between the DMN and SN is greater following LC-NE activation under our experimental condition. This finding may suggest a putative role of tonic LC-NE activity in improving efficient cognitive control and reducing behavioral variability by fostering SN-DMN anticorrelation (*72*). Additionally, our findings point to the possibility that tonic and phasic outputs from LC-NE neurons may preferentially drive DMN and SN, respectively. Further, the enhanced anticorrelation between networks also supports the plausible mechanism by which NE-targeting pharmacological agents like atomoxetine, clonidine, and guanfacine are effective in treating Attention-Deficit/Hyperactivity Disorder (ADHD) patients since they strengthen the FC within the *Frontal* DMN module and result in improved attention (*73*).

Our DCM and mediation analyses show that enhanced *Frontal* module connectivity in the DMN was causally manipulated by reducing *RSC-HIPP* control of the *Association* module, with the connectivity between the *Association* and *Frontal* modules serving as a key mediator. These findings reveal a new understanding of how LC-NE activation controls the signaling cascades within DMN modules and achieves its control of the *Frontal* cortical regions, which are among the most well-studied projection targets of LC-NE due to their importance in shaping multiple behaviors (*74*). LC-NE neuronal loss and the subsequent depletion of cortical NE levels are widely considered to be among the first sites of neurodegeneration in Parkinson’s and Alzheimer’s disease (*75*), resulting in behavioral pathologies linked to alterations in DMN such as delayed attention shifting (*76*), enhanced mind-wandering (*77*), and reduced cognitive and emotional processing of sensory information (*76, 77*). Optogenetic-induced LC-NE activation at lower tonic frequencies into prefrontal and orbitofrontal cortices enhances stimulus and goal-directed attention with decreased impulsivity (*78*). Conversely, the suppression of LC-NE activation exacerbates distractibility and impulsivity (*78*), similar to that observed in Parkinson’s and Alzheimer’s disease patients (*77*). Taken together, alterations within *Frontal* and between *Frontal* and *RSC-HIPP* DMN modules have potential to serve as early biomarkers for pathophysiological changes in LC-NE neurons. Our circuit-level findings could also pave the way towards novel targets to causally control the *Frontal* DMN via the *RSC-HIPP* module when LC neurons have degenerated, such that the behavioral traits relevant to the *Frontal* DMN may be restored when endogenous NE is pathologically diminished.

## MATERIAL AND METHODS

### Animals

All animal procedures were performed in strict compliance with ethical regulations for animal research and approved by the Institutional Animal Care and Use Committee of the University of North Carolina at Chapel Hill. *En1*^*cre*^ (*79*), *Dbh*^*Flpo*^ (*31*) and *RC::FL-hM3Dq* (*30*) mouse colonies are maintained on a C57BL/6J background. Male and female triple transgenic animals were generated at the National Institute of Environmental Health Sciences by crossing *En1*^*cre*^ mice to double transgenic *Dbh*^*Flpo*^; *RC::FL-hM3Dq* mice. Single- and double-transgenic littermates served as controls. All animals were maintained on a 12/12 h light-dark cycle with access to food and water *ad libitum*.

### CBV-fMRI acquisition

For fMRI studies, LC-NE (n=9) and control mice (n=12) were initially anesthetized using 2-3% isoflurane and maintained under light anesthesia (1% isoflurane) while preserving physiological homeostasis. All MRI experiments were performed on a Bruker BioSpec 9.4-Tesla, 30 cm bore system (Bruker BioSpin Corp., Billerica, MA) with ParaVision 6.0.1 on an AVANCE II console. An RRI BFG 150/90 gradient insert (Resonance Research Inc., Billerica, MA) paired with a Copley C700 gradient amplifier (Copley Controls Corp., Canton, MA) were used for all experiments. A 72 mm volume coil was used as the transmitter and quadrature mouse brain coil was used as the receiver (Bruker BioSpin Corp., Billerica, MA). Magnetic field homogeneity was optimized first by global shimming, followed by local second-order shims using a MAPSHIM protocol. All CBV-fMRI data were acquired using a 2D multi-slice, single-shot, gradient-echo echo-planar imaging sequence: TR (repetition time) = 3000 ms, TE (echo time) = 7.9 ms, bandwidth = 250 kHz, flip angle = 70 degrees, FOV (field of view) = 19.2 × 19.2 mm^2^, matrix size = 64 × 64, slice number =26, slice thickness =0.3 mm, resulting in isotropic voxel size of 0.3 mm^3^. Subjects were continuously scanned for 40 min with a 1 mg/kg dose of CNO administered via an intraperitoneal catheter 10 min after scan onset. CBV-weighted fMRI was achieved by a bolus dose of an in-house developed iron oxide nanoparticles (30 mg Fe/kg, i.v.)(*80*). Rectal body temperatures were continuously maintained at 37±0.5°C by a temperature controller (Oakton Temp9500, Cole-Parmer, Vernon Hills, IL, USA) coupled to a circulating water bath (Haake S13, Thermo Scientific, Waltham, MA, USA) that heats the MRI mouse cradle. Respiration was monitored through a pneumatic pillow (Respiration/EEG Monitor, SA Instruments, Stony Brook, NY, USA) and maintained between 90-110 breaths/min.

### fMRI data analysis

#### Pre-processing of images

All fMRI data were corrected for slice timing and motion using AFNI. Brain data were isolated using a U-Net deep-learning skull stripping tool and spatially normalized to our EPI template using ANTs. Additionally, de-spiking, ICA de-noising, and nuisance variable regression of the six motion parameters estimated from motion correction and the CSF signal extracted using a mask of the major brain ventricles. Datasets were then smoothed using a Gaussian kernel with full width half maximum (FWHM) at 0.6 mm, de-trended, and temporally filtered by applying a high-pass filter between > 0.01 Hz. Datasets underwent quality control by measuring frame wise displacement (FD), temporal SNR (tSNR), and DVARS (temporal derivative of the root mean squared variance over voxels). A detailed description of the preprocessing pipeline (*55*) can be found in the **Supplementary Methods**.

#### ICA and modularity analyses

MRI data were decomposed into 100 functional components using baseline data from all subjects via a group-level ICA (FSL MELODIC). Functional modules of the identified 17 DMN components were parcellated using the Louvain community detection algorithm. Within- and between-module connectivity was then defined as the average of FC across node pairs within or between the identified DMN modules. The ICA and modularity analysis method is detailed in the **Supplementary Methods**.

#### DCM and mediation analysis

We specified a DCM model with full connectivity consisting of three modules from DMN to estimate pairwise EC among the DMN modules and constructed a directed and weighted graph (representing an EC network) for each subject. We applied serial-multiple mediation analysis model in AMOS 17.0 (SPSS Inc., Chicago, IL, USA) to uncover underlying functional pathways within DMN. Specifically, we first estimated the direct relationships between dependent variable (EC from *RSC-HIPP* module to *Association* module) and independent variable (FC within *Frontal* module). Then, in a mediation model, the FC between *Association* and *Frontal* module was added as a mediator. In this context, full mediation occurs when the relationship between the independent variable and the dependent variable is no longer significant with the inclusion of a mediator variable. Detailed data analysis is further described in the **Supplementary Methods**.

### FDG PET procedure

Mice were fasted 12 h prior to undergoing ^18^F-FDG PET scans to reduce variability in blood glucose levels that could alter ^18^F-FDG uptake (117). Static PET scans were collected on the same cohort of animal over two scan sessions using LC-NE/hM3Dq (n=5) and control (n=6) mice to represent sham-treated baseline or CNO-treated condition. Mice were briefly anesthetized under 1-3% isoflurane and injected with either a saline + DMSO vehicle or CNO dissolved in DMSO (1 mg/kg, i.p.) and subsequently received an i.v. injection of ∼0.2 mCi of ^18^F-FDG after 5 min. Mice were recovered in their home cages for a 45 min uptake period. Mice subsequently were anesthetized with isoflurane (2%) and underwent a 10 min CT and 20 min PET scan on a small animal PET/CT scanner (Argus-2R, Sedecal, Madrid, Spain). PET data were reconstructed using the 2D ordered subset expectation maximization (OSEM) algorithm expressed in standardized uptake values (SUV) and normalized using arm muscle uptake of ^18^F-FDG using PMOD (PMOD Technologies LLC, Zurich, Switzerland). Data were represented as % changes in SUV between vehicle and CNO scans.

### Fiber-photometry procedure

#### Surgical AAV microinjection and fiber implantation

LC-NE (n=5) and control (n=5) mice were microinjected 0.3 µl of AAV5-hsyn-NE_2.1_ (h-N01, WZ Biosciences) and 0.5 µl of AAV9-hsyn-jRGECO1a (100854, Addgene) to the left Cg1 (AP=2.2mm, ML=0.2mm, DV=-1.6mm). NE_2.1_ (a green fluorescent norepinephrine sensor) and jRGECO1a (a red-shifted intracellular calcium sensor) were used for determining the NE release and neuronal activity, respectively. An optic fiber was implanted 0.3 mm above the injection site and imbedded to the skull using cement (C&B Metabond, S380, Parkell).

#### Fiber-photometry recording

All recordings began at least four weeks after surgery. CY5-conjugated Dextran fluorescent dye (20 mg/kg, R-FN-006, RuixiBio) was injected intravenously for CBV measurements. Animals were prepared and maintained under the same conditions as fMRI experiments. A spectral fiber-photometry system capable of recording NE_2.1_, jRGECO1a, and CY5 signals were simultaneously recorded during the experiments. Detailed methods can be found in the **Supplementary Methods**.

### Immunohistology procedure

Mice were deeply anesthetized with sodium pentobarbital and transcardially perfused with 0.1 M phosphate buffered saline (PBS) followed by 4% PFA. Brains were post-fixed overnight by immersion in 4% PFA at 4°C. Following a rinse in PBS, brains were cryoprotected in 30% sucrose in PBS and embedded in Tissue Freezing Medium. 40 µm free-floating coronal cryosections were collected in PBS and processed for immunohistochemistry according to previously published protocol (*31*). In brief, free-floating sections were blocked in 5% normal goat serum in PBS with 0.1 % Triton X-100) for 1 h prior to incubating in primary antibody overnight at 4°C. Primary antibodies against mCherry (EST202, Kerafast), GFP (AB13970, Abcam) and TH (AB152, Millipore). Sections were washed in PBS 3x and incubated for 2 h in the secondary antibody the following day. Secondary antibodies were Alexa Fluor 488, 568, 648 (Invitrogen). Sections were mounted onto glass slides, coverslipped with Vectashield hard-set mounting medium with DAPI (H-1500, Vector Labs) and imaged on a Zeiss LSM 880 inverted confocal microscope.

## Data availability

All MRI, PET, and photometry data from this study are openly available on Mendeley Data (https://doi.org/10.17632/hxch8htz84.1).

## Acknowledgements

This research was supported by the Extramural Research Programs of US National Institutes of Health, NINDS (R01NS091236), NIMH (R01MH126518, R01MH111429, RF1MH117053), NIAAA (P60AA011605 and U01AA020023), and NICHD (P50HD103573) to Y.Y.I.S and the Intramural Research Program of the US National Institutes of Health, National Institute of Environmental Health Sciences (ZIA-ES102805 for P.J. and 1ZIAES103310 to for G.C.). We thank the members of the UNC Center for Animal MRI and Drs. Fulton Crews, Zoe McElligott, and Bryan Roth for their inputs. We thank UNC Small Animal Imaging Core Facility staff Jon Frank and Joseph Merrill for their assistance in PET data acquisition.

## Author contributions

E.A.O., M.D., and Y.I.S. designed the study. E.A.O and M.D. collected the imaging data. E.A.O., L.M.H and S.L. analyzed the imaging data. J.Z. and G.C. performed the viral injections and fiber implantation. E.A.O., T.H.C. and W.Z. collected and analyzed the fiber photometry data. K.S. processed and K.S. and P.J. analyzed the histology data. N.S., I.E. and P.J. developed and shared the transgenic mice. E.A.O., L.M.H., P.J. and Y.I.S. wrote the manuscript with input from all authors.

## Additional information

Supplementary methods and figures accompany this paper.

