## Supplementary Materials for "Chemogenetic Stimulation of Tonic Locus Coeruleus Activity Strengthens the Default Mode Network"

This PDF file includes:

**Supplementary Methods**
**Supplementary Figures**
**Fig. S1.** LC-NE neurons send projection into DMN brain regions
**Fig. S2.** Representative images of raw fMRI acquisitions and fMRI atlas templates
**Fig. S3.** 100 component ICA of the mouse brain with quality assurance data
**Fig. S4.** Mouse DMN modules and interconnectedness quality assurance data
**Fig. S5.** Detailed CBV, ReHo, ALFF data
**Fig. S6.** Validation of circulating far-red fluorescent CBV dye decay
**Fig. S7.** Detailed PET data
**Fig. S8.** DMN graph properties and FC association with insula
**Fig. S9.** DCM of controls
**Fig. S10** Modulation analysis of controls

### Supplementary Methods

#### Axonal Labeling of Transgenic Mice

To label LC-NE axons, *En1*$^{cre}$ mice were crossed to mice expressing *Dbh*$^{Flpo}$ and the dual recombinase reporter *RC::FLTG* (1) to generate *En1*$^{cre}$; *Dbh*$^{Flpo}$; *RC::FLTG* mice that express GFP throughout LC-NE neuronal projections. Brains were post-fixed overnight by immersion in 4% PFA at 4°C. Following a rinse in PBS, brains were cryoprotected in 30% sucrose in PBS and embedded in Tissue Freezing Medium. 40 µm free-floating coronal cryosections were collected in PBS and incubated with primary antibodies for GFP (AB13970, Abcam) and NET (1447-NET, Phosphosolutions). Secondary antibodies were Alexa Fluor 488 and 633 (Invitrogen). After washing, the sections were stained with Neurotrace 435/455 blue fluorescent Nissl stain (1:50, N21479, Thermo Fisher) and mounted using Prolong Diamond anti-fade mountant (Thermo Fisher, Scientific, Waltham, MA, USA). Sections were imaged on a Zeiss LSM 880 inverted confocal microscope with a 40x objective. Zen Black Software (Carl Zeiss) was used to convert z-stacks to maximum intensity projections. Images were modified only by adjusting brightness and contrast across the entire image to optimize the fluorescence signal.

#### fMRI Pre-processing

*Slice-Time Correction*: Slice-time correction interpolating the temporal offsets during interleaved acquisitions of slices to eliminate radio frequency pulse excitation leakage artifacts. Correction was done by adjusting slice timing to a reference time point using 3dTshift in AFNI.

*Motion Correction:* Motion correction applies rigid-body transformation to align all temporal volumes to a reference volume to correct for head motion during fMRI acquisitions that may produce spurious correlations if uncorrected. Our pipeline utilizes 3dvolreg in AFNI.

*Brain Segmentation:* CBV-fMRI time series were averaged to single volume and corrected for image intensity non-uniformities using N4BiasFieldCorrection in ANTs. Brain segmentation was performed on averaged, bias-corrected imaged using an established U-Net deep-learning skull stripping tool (2) and applied across the time series.

*Spatial Normalization*: We first generated an averaged interim CBV-fMRI mouse atlas which was co-registered and placed through linear and non-linear warping using antsRegistrationSyN.sh in ANTs into the template space of the Allen Mouse Brain Atlas (v3) (3). The Allen Mouse Brain Atlas utilizes the Mouse Common Coordinate Framework allowing researchers to overlap their data into the standardize template space. We then spatially normalized all individual subject (n=21) to our interim CBV-fMRI mouse atlas on ANTs. Linear affine transformation matrices and warp field maps were applied across the entire CBV-fMRI time series using WarpTimeSeriesImageMultiTransform in ANTs. A final CBV-fMRI mouse atlas was generated for future studies using all spatially normalized subjects into the Allen Mouse Brain Atlas template space using the deformation template generator antsMultivariateTemplateConstruction2.sh in ANTs to form averaged atlases that are openly available in the posted study data.

*Cleaning and De-noising fcMRI data:* Spatially normalized fcMRI datasets were first de-spiked using 3dDespike in AFNI to eliminate large signal intensity spikes that surpassed ±2.5 standard deviations above the local median absolute deviation calculated at every time frame against ±4 adjacent time frames. Next, the data underwent nuisance variable regression based on a General Linear Model (GLM) using 3dDeconvolve in AFNI to eliminate the contribution of measurable noise from non-meaningful sources such as head motion and periodic physiological pulsations that include cardiac, respiratory, vascular and CSF oscillations. The GLM included six motion regressors (accounting for motion correction in the X, Y and Z translation directions and pitch, yaw, and roll in rotation directions) and CSF oscillations extracted from a mask layer of the ventricles of the mouse brain to routinely account for cardiac pulsation noise. We did not regress the global signal since it has been demonstrated to introduce spurious anti-correlations throughout the brain in some cases (4). The data then underwent spatial smoothing using 0.6 mm full width at half maximum (FWHM) Gaussian kernel to improve signal to noise ratio (SNR). The data was then de-trended to remove drifting likely associated with MRI gradient heating during acquisitions. Lastly, the data underwent temporal filtering using a bandpass filter for frequencies between 0.01-0.01 Hz to extract meaningful low-frequency oscillations while eliminating high frequency noises/artifacts.

**fMRI Analysis**

*CBV Calculations:* We calculated the percentage change in CBV using the following equation:

$$\% \, CBV = \frac{\frac{-1}{TE} \ln \left(\frac{S_{CNO}}{S_{preCNO}}\right)}{\frac{-1}{TE} \ln \left(\frac{S_{preCNO}}{S_0}\right)}$$

where $S_0$ represents MR signal intensity before iron oxide administration, $S_{preCNO}$ represents MR signal intensity after iron oxide administration but before CNO administration, and $S_{CNO}$ represents MR signal intensities after CNO administration.

*ReHo and ALFF analyses of DMN modules:* ReHo (5) is a voxelwise analysis to determine local synchronization of fMRI signal among all brain regions. Higher ReHo scores typically represent greater neuronal coherence and centrality within nodes but does not necessarily equate to higher neuronal activity. ALFF (6) is a voxelwise technique to calculate the total power of regional neuronal activity within 0.01-0.1 Hz. Overlapping regions that share high ALFF and ReHo scores are thought to have enhanced synchronization of low frequency neuronal activity.

*ICA analysis:* MRI data were decomposed into 100 FCs using baseline data from all subjects via a group-level ICA (FSL MELODIC) (7). ICA spatial maps were back-reconstructed for each subject to estimate subject-specific temporal components and associated spatial maps using dual-regression analysis (8), and a one-sample two-sided t-test was performed to generate group component maps. Individual time courses from 17 pre-selected DMN nodes were extracted from the first stage of dual-regression and the Fisher z-transformed Pearson correlation were computed among all node pairs to form a correlation matrix. To keep the strongest connections without isolated nodes, the z-threshold = 0.26 (connection density = 48%) was set in the mean FC matrix across all subjects. Functional modules of the DMN were parcellated using the Louvain community detection algorithm (9).

*Modularity Analysis:* Modules refer to groups of nodes that are highly connected with each other but less connected with other nodes in a network. The modularity $Q$ quantifies the efficacy of partitioning a network into modules by evaluating the difference between the actual number of intramodule connections and the expected number for the same modules in a randomized network. The objective of a module detection procedure is to find a specific partition that maximizes the modularity $Q$. Newman's spectral algorithm was used for modular detection. To examine whether a network had significantly higher modularity than the random graphs, we randomized the original network with preserved strength distribution 1,000 times and calculated the mean ($\mu$) and standard deviation ($\sigma$) of those modularity values. We compared the modularity $Q$ of the real network to those values:

$$Z = \frac{Q - \mu}{\sigma}$$

which measures how many standard deviations the real modularity is above the mean for the random graph.

*Sobel Test Calculations:* The mediation effect of each significant path was evaluated using the Sobel test whereby the z-value was estimated using the following equation:

$$z - value \ = \frac{a \times b}{\sqrt{b^2 \times S_a{}^2 + a^2 \times S_b{}^2}},$$

where $a$ is the mean regression coefficient for the association between independent variable and mediator, $b$ is the mean regression coefficient for the association between mediator and dependent variable, $S_a$ is standard error of $a$, and $S_b$ is standard error of $b$.

*FC analysis within and between DMN modules:* Within- and between-module connectivity was defined as the average of FC across node pairs within or between the identified DMN modules. We conducted network-based statistics (NBS) (10) to investigate the significant connection changes. For each comparison, a primary component-forming threshold (*p*<0.05, uncorrected) was applied to form a set of supra-threshold edges and all the remaining connected subnetworks in the matrix were then evaluated under the null hypothesis of random group membership (5,000 permutations).

*DCM analysis:* We specified a DCM model (11) with full connectivity consisting of three modules from DMN. After extracting the fMRI time series from the DMN modules through dual-regression analysis, we used spectral DCM in SPM12 (12) to estimate pairwise EC among the DMN modules and constructed a directed and weighted graph (representing an EC network) for each subject.

*Mediation analysis:* We applied serial-multiple mediation analysis model using the structural equation modeling method (13) in AMOS 17.0 (SPSS Inc., Chicago, IL, USA) to uncover underlying functional pathways within DMN. Specifically, we first estimated the direct relationships between the dependent variable (EC from *RSC-HIPP* module to *Association* module) and the independent variable (FC within *Frontal* module). Then, in a mediation model, the FC between the *Association* and *Frontal* modules was added as a mediator. In this context, full mediation occurs when the relationship between the independent variable and the dependent variable is no longer significant with the inclusion of a mediator variable (14). For each pathway, we utilized the bootstrapping test 1,000 times to test the significance. Finally, the mediation effect was evaluated using the Sobel test (15).

**Fiber Photometry**

The fiber photometry setup for this study required triple-excitation continuous wave lasers at 488 nm (OBIS Galaxy 488nm LX 100mW, 1236444, Coherent Inc., Santa Clara, CA, USA), 561 nm (OBIS Galaxy 488nm LS 80mW, 1275608, Coherent Inc., Santa Clara, CA, USA) and a 644 nm (OBIS Galaxy 640nm LX 75mW, 1236445, Coherent Inc., Santa Clara, CA, USA) housed in an OBIS LX/LS Laser Box (1228877, Coherent, Inc.). The lasers were aligned and combined using OBIS Galaxy Laser Beam Combiner (1253556, Coherent Inc., Santa Clara, CA, USA) and the beam was sent through a neutral density filter (NEK01, Thorlabs, Newton, NJ) and a dichroic mirror (ZT405/488/561/640rpcv2, Chroma Technology Corp., Bellow Falls, VA, USA) within a fluorescence cube (DFM1, Thorlabs, Newton, NJ, USA). The combined laser beam was cast through an achromatic fiber port (PAFA-X-4-A, Thorlabs, Newton, NJ, USA) into a multimode optical fiber patch cable with a 105 µm core (M61L01, Thorlabs, Newton, NJ, USA), terminating into a 1.25 mm outer diameter ceramic ferrule connected to the surgically implanted optical fiber cannula via a ceramic spilt sleeve (SM-CS125S, Precision Fiber Products Inc., San Diego, CA, USA). Emitted fluorescent signals travel back along the patch cable, through the emission filters of the dichroic mirror and emission filter (ZET405/488/561/640mv2, Chroma Technology Corp., Bellow Falls, VA, USA), then launch through an aspheric fiber port (PAF-SMA-11-A, Thorlabs, Newton, NJ, USA) into the core of an AR-coated 200/230 mm core/cladding multi-mode patch cable (M200L02S-A, Thorlabs, Newton, NJ, USA). The AR-coated multi-mode patch cable was connected to a spectrometer (QE Pro-FL, Ocean Optics, Largo, FL, USA) for spectral data acquisition, which was operated by a UI software OceanView (Ocean Optics, Largo, FL, USA).

In order to perform spectral linear unmixing of the spectroscopy data, we used the following linear regression algorithm:

$$Y(t) = A + Co_{NE2.1} * S_{NE2.1} + Co_{jRGECO1a} * S_{jRGECO1a} + Co_{CY5-Dextran} * S_{CY5-Dextran} + \varepsilon(t)$$

whereby $Y(t)$ is the observed mixed spectrum at any given time and $S_{NE2.1}$, $S_{jRGECO1a}$ and $S_{CY5-Dextran}$ are the normalized reference emission spectra for $NE_{2.1}$, jRGECO1a and CY5-conjugated Dextran fluorescent dye, respectively. $A$ is an unknown constant while $Co_{NE2.1}$, $Co_{jRGECO1a}$, and $Co_{CY5-Dextran}$ are unknown regression coefficients that correspond to the green, red, and far-red channels, respectively. Lastly, $\varepsilon(t)$ denotes the random error associated with the model. The model allows us to estimate $A$, $Co_{NE2.1}$, $Co_{jRGECO1a}$, and $Co_{CY5-Dextran}$ at each time point. The $Co_{NE2.1}$, $Co_{jRGECO1a}$, and $Co_{CY5-Dextran}$ were further detrended according to their baseline period using a homemade Matlab script. The spike SNR during the first 10 min baseline and the last 10 min post-CNO recording were calculated following high-pass filtering (cut at 0.1 Hz) on z-transformed time segments. Any local maximum with z-score > 1.96 was considered a spike.

### Supplementary Figures

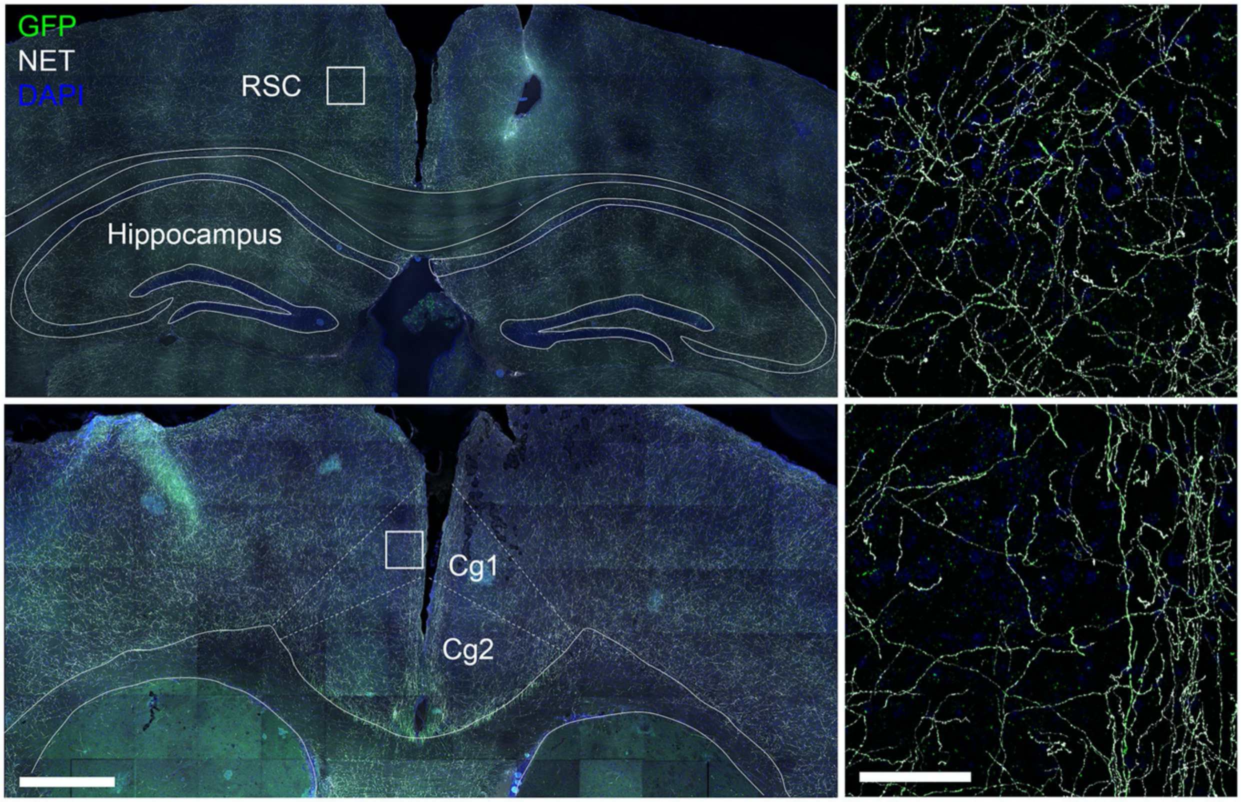

**fig. S1** LC-NE projections to DMN brain regions. To label LC-NE axons, *En1*$^{cre}$ mice were crossed to mice expressing *Dbh*$^{Flpo}$ and the Flp/Cre-responsive recombinase reporter *RC::FLTG* . Recombination of *RC::FLTG* by *En1*$^{cre}$ and *Dbh*$^{Flpo}$ results in GFP expression in LC-NE neurons. Coronal sections show innervation of LC-NE axons (green) co-localized with the noradrenergic transporter, NET (white) in the Cg1 and RSC of the DMN.

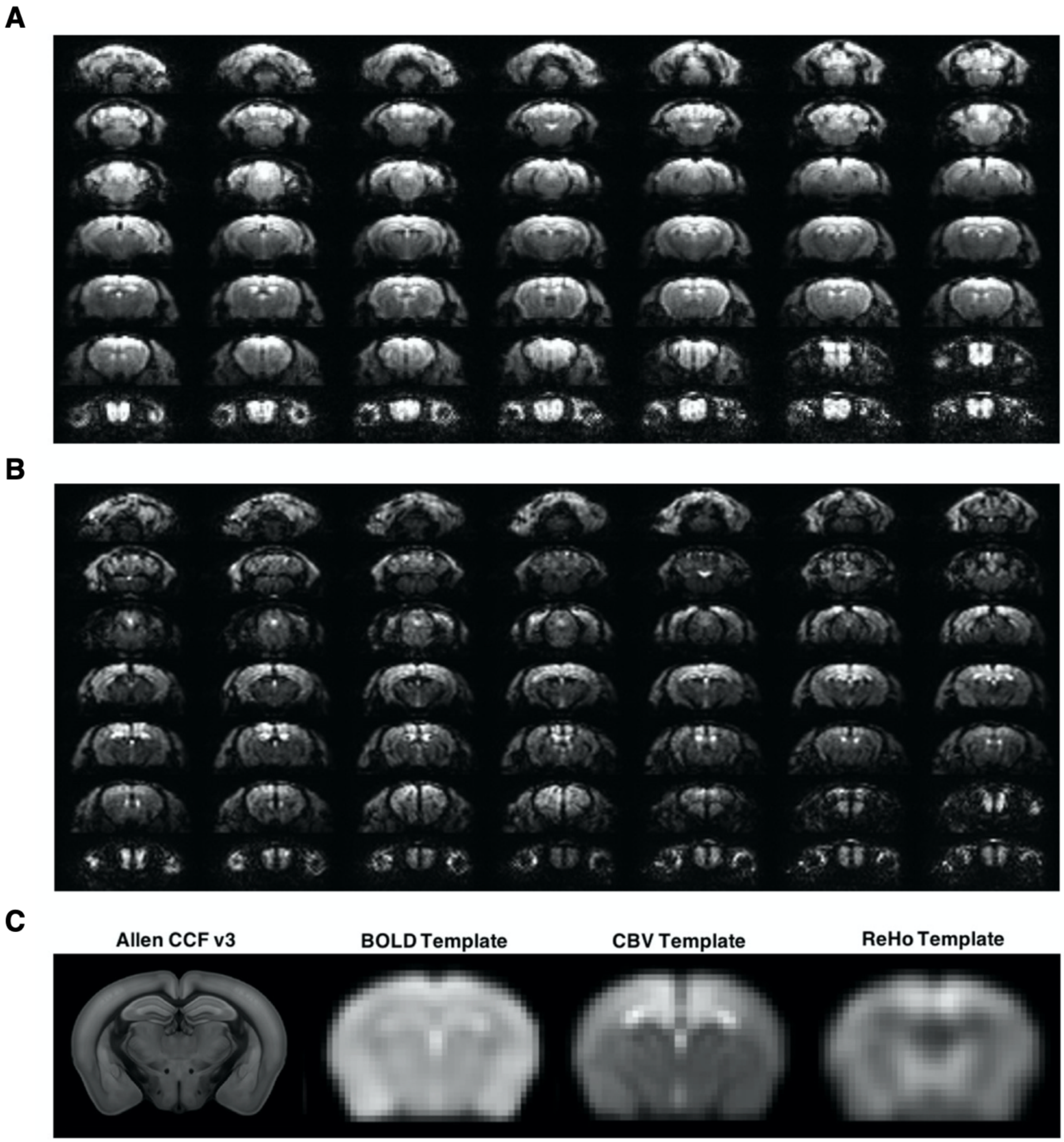

**fig. S2** Representative single-shot gradient echo planar images. **(A)** BOLD-weighted contrast. **(B)** CBV-weighted contrast. **(C)** EPI scans (n=21) were spatially warped to the Allen Mouse Common Coordinate Framework and Reference Atlas (v3) using to generate symmetric diffeomorphic atlases for BOLD, CBV and ReHo.

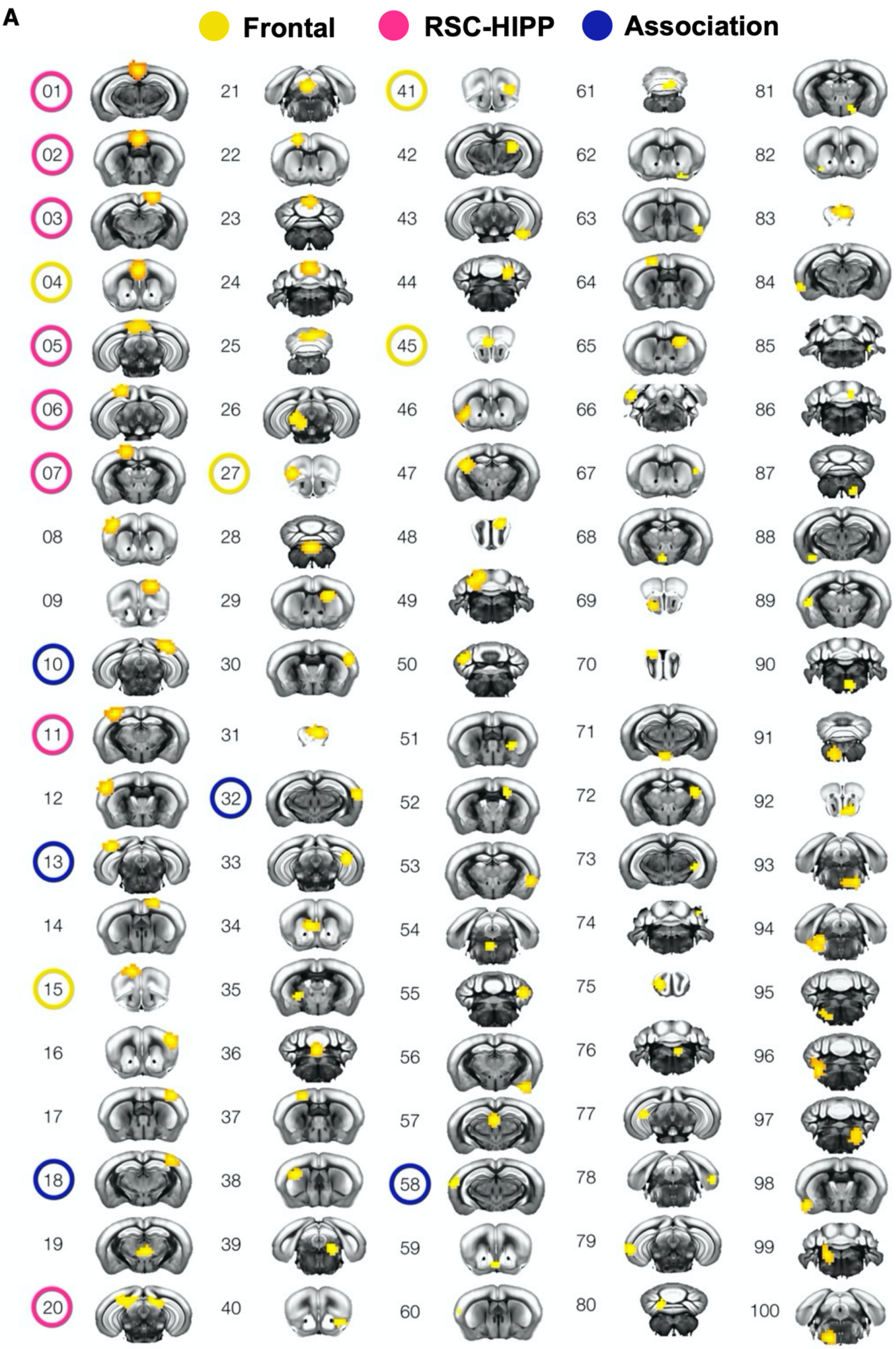
A
Frontal
RSC-HIPP
Association

**B**

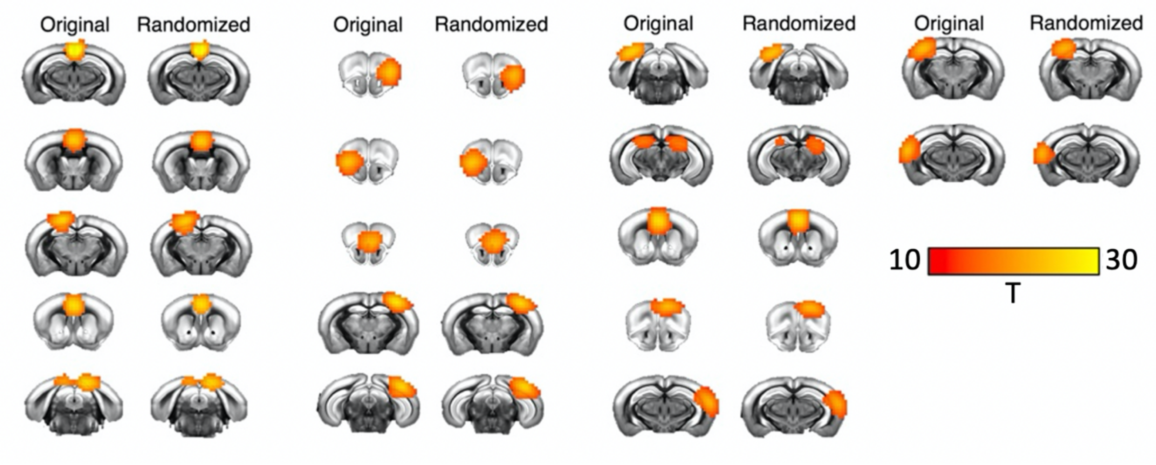

**fig. S3** Detailed ICA maps from baseline fMRI scans. **(A)** Group ICA map of 100 ICs derived from baseline fMRI of all subjects (n=21) prior to CNO administration. **(B)** To assure the reproducibility of components, ICA was conducted again using randomly order components. Highly similar components could be found in the randomized data. T-value threshold = 10.

**A**

| IC | Approximate Brain Region |
|---|---|
| 1 | Retrosplenial Granular Cortex (RSG) |
| 2 | Anterior Cingulate Cortex 2 (a-Cg2) |
| 3 | right Medial Parietal Association Cortex (R-MPtA) |
| 4 | anterior Cingulate Cortex 2 (aCg1) |
| 5 | posterior Cingulate Cortex 2 (p-Cg2) |
| 6 | left Retrosplenial Dysgranual Cortex (L-RSD) / Visual Cortex (L VIS) |
| 7 | left Medial Parietal Association Cortex (L MPtA) |
| 10 | right RSD / Visual Cortex (VIS) |
| 11 | left Posterior Parietal Cortex (PPtC) |
| 13 | left Visual Cortex (L VIS) / Posterior Parietal Cortex (L PPtC) |
| 15 | Prelimbic (PrL) & Infralimbic (IL) Cortices |
| 18 | right Posterior Parietal Cortex (R PPtC) |
| 20 | dorsal Hippocampus (HIPP) |
| 27 | left Lateral Orbital Cortex (L LO) |
| 32 | right Auditory (R Aud) & Temporal Association (R TeA) Cortices |
| 41 | right Lateral Orbital Cortex (R LO) |
| 45 | Cingulate Cortex 1 (Cg1) |

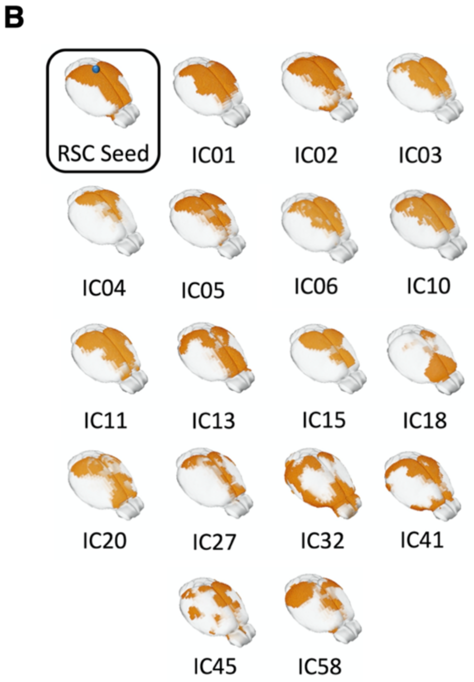

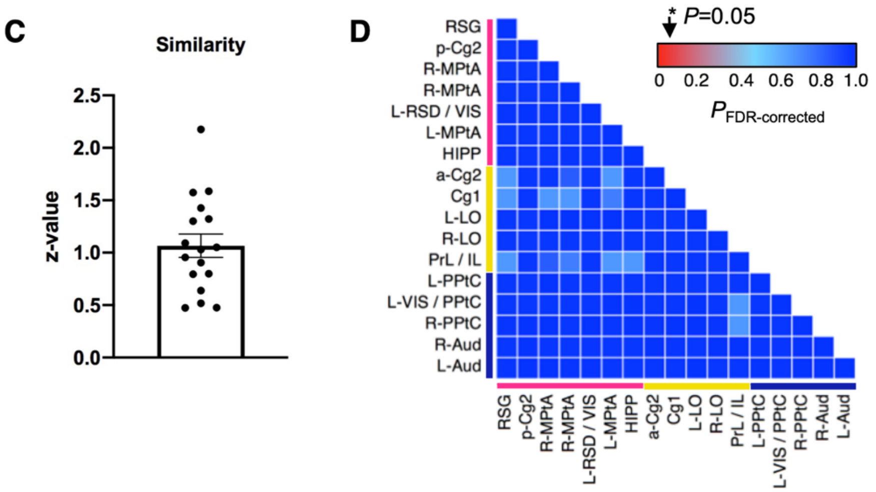

**fig. S4** Detailed evaluation of ICs associated with DMN. **(A)** Selected DMN ICs and their corresponding DMN brain regions. **(B)** Dual regression results of the 17 DMN ICs and RSC seed-based connectivity map ($P$<0.001). **(C)** Spatial similarity between the RSC seed-based connectivity map and the dual regression results of all 17 DMN ICs (One-sample t-test, $P$<0.01). **(D)** To access the baseline variability, ANOVA was performed among the baseline scans of LC-NE and control groups. No significant FC change was identified between subject groups across 17 ICs.

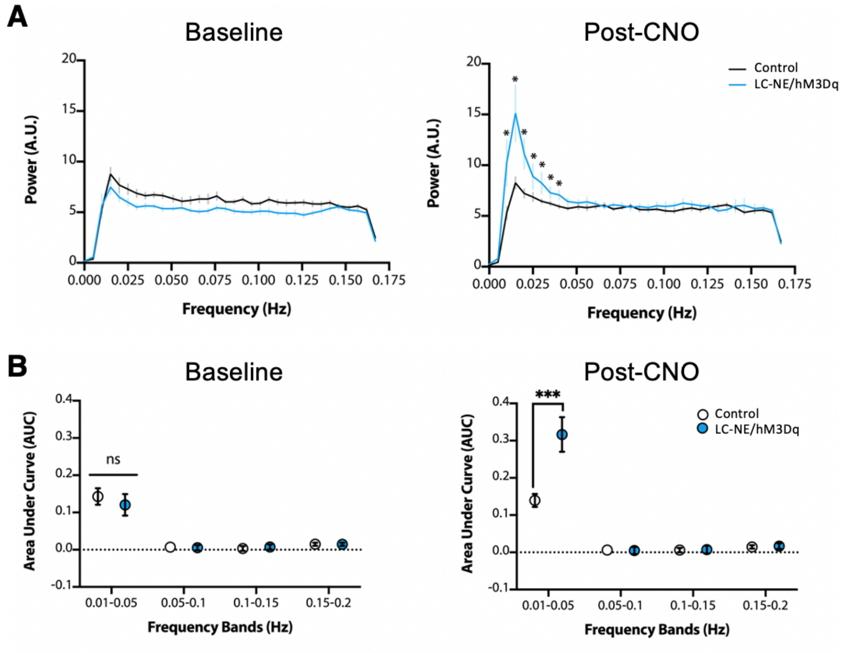

**fig. S5** ALFF changes following LC-NE activation. **(A)** Periodogram of ALFF spectral signals in the *Frontal* module shows significant increase in power between 0.01-0.05 Hz following LC-NE activation. **(B)** Area-under-the-curve of ALFF changes following LC-NE activation were restricted to frequency bands ranging from 0.01-0.05 Hz. *$p < 0.05$, *** $p < 0.001$, error bars represent ± SD.

$$Y = 16.79 + 48.24^{(-0.09072 * X)} + 34.70^{(-0.01065 * X)}$$

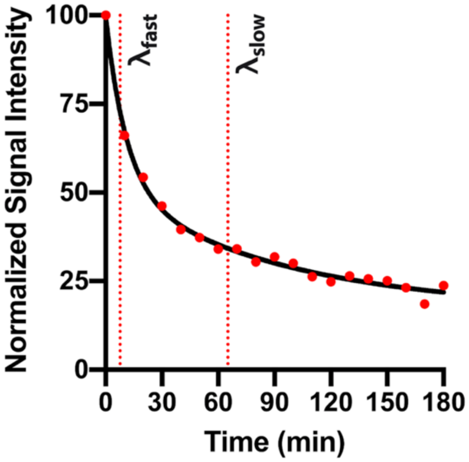

| Two-Phase Decay | |
|---|---|
| $Y_0$ | 99.73 |
| Plateau | 16.79 |
| Percent Fast | 58.16 |
| $K_{Fast}$ | 0.091 |
| $K_{Slow}$ | 0.011 |
| $\lambda_{Slow}$ | 65.07 |
| $\lambda_{Fast}$ | 7.64 |
| $\tau_{Slow}$ | 93.88 |
| $\tau_{Fast}$ | 11.02 |
| Rate Constant Ratio | 8.517 |

**fig. S6** Wash-out of CY5-Dextran dye in the bloodstream. High goodness-of-fit ($R^2$=0.9937) to a two-phase exponential decay curve was observed, likely due to rapid clearance of unconjugated dye during the fast phase and a slower wash-out phase of conjugated dye. All photometry data reported in this study were collected during the second wash-out phase where the trend is closer to linear. The trend was further removed by fitting a linear slope during pre-CNO baseline period.

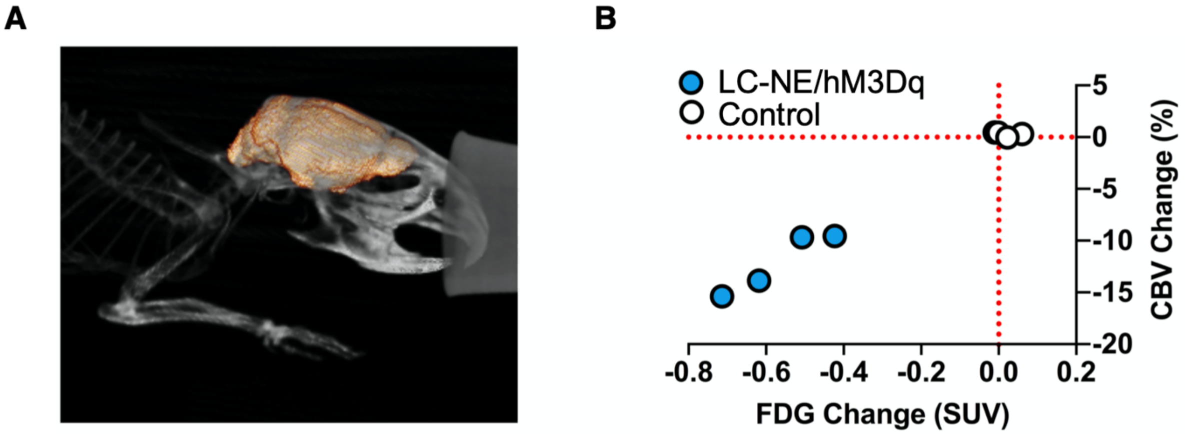

**fig. S7** Comparison of FDG uptake and changes in CBV. **(A)** A representative 3D rendered brain mask for PET analysis. Skull-stripping was guided by the CT scan of the same subject. **(B)** CBV and FDG uptake changes in the *Frontal* DMN module among LC-NE/hM3Dq and control subjects that underwent both modalities. A robust correlation was observed (r=0.9218, $p<0.05$), suggesting a relationship between FDG and CBV observations following the activation of LC-NE neurons.

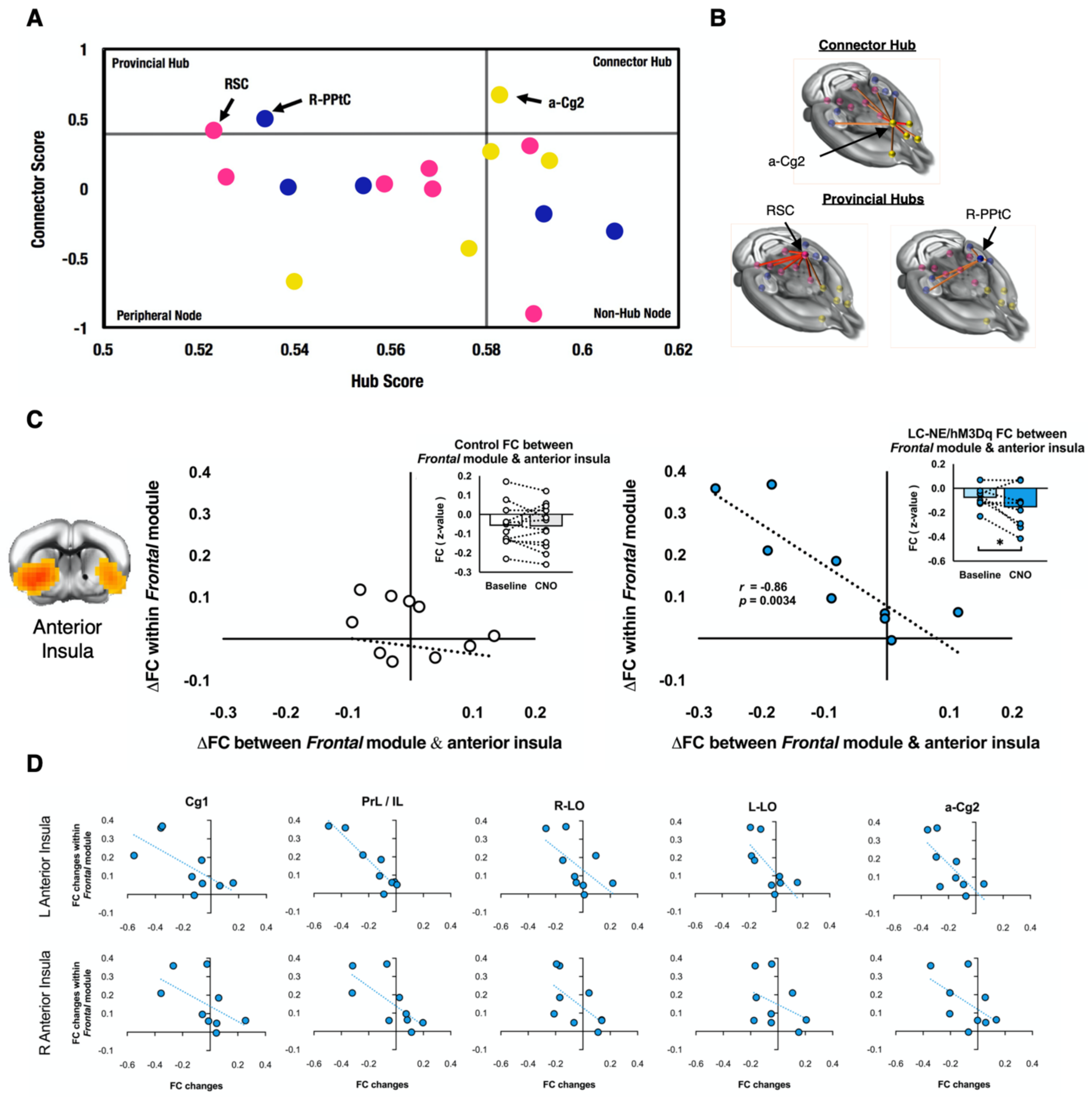

**fig. S8** Graph properties of the mouse DMN nodes and anti-correlations between DMN and anterior insula following LC-NE activation. **(A)** Participation coefficient and within-module degree were estimated to identify hubs and connectors in the DMN, respectively. **(B)** Hubs and their respective FCs to other nodes. **(C)** Comparisons of FC changes within *Frontal* module against FC changes between the *Frontal* module and anterior insula in LC-NE mice. The observed negative correlation suggests the existence of antagonizing relationship (r=-0.86, $p$<0.005). **(D)** *Post-hoc* analyses of the relationships between FC changes within the *Frontal* module and FC changes between the anterior insula and each *Frontal* module node in LC-NE/hM3Dq mice. * $p$-values are FDR-corrected.

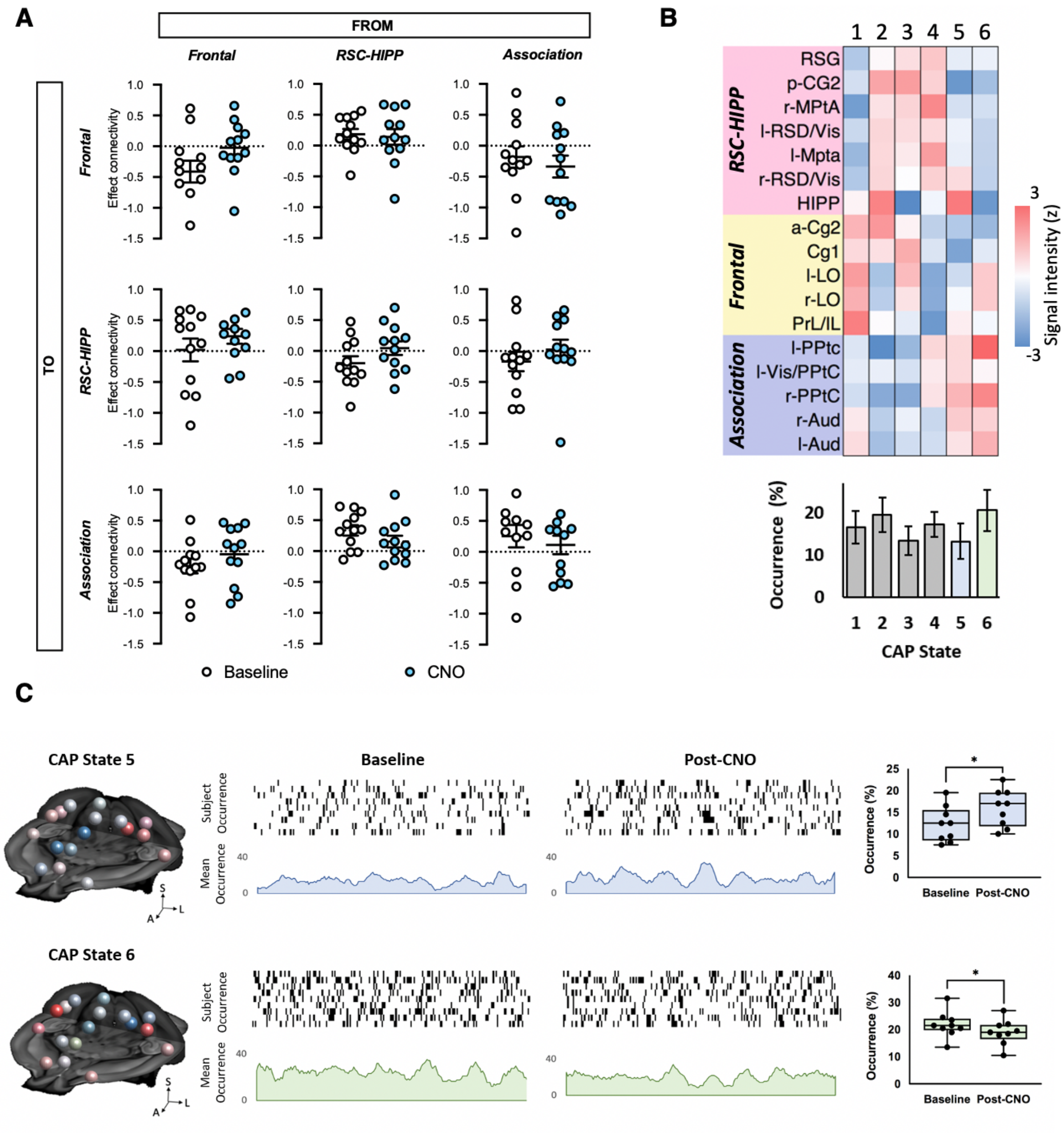

**fig. S9 (A)** Dynamic causal modeling (DCM) analysis among DMN modules of the littermate control groups. No causal associations were detected following CNO injection. Horizontal lines represent means, and error bars represent ± SD. **(B)** k-means clustering identified six CAP states among the DMN nodes. **(C)** LC-NE significantly increased the occurrence of CAP state 5 and significantly decreased the occurrence of CAP state 6. No changes were found in other CAPs. State 5 represents isolated HIPP activation that is asynchronous with the rest of the *RSC-HIPP* module nodes, whereas State 6 represents synchronous *RSC-HIPP* and *Association* modules with opposing polarities. While CAP data do not infer causality, the high occurrence of State 6 suggests a robust state recapitulating that *RSC-HIPP* and *Association* activity are opposed during resting state, but become less polarized following CNO administration. *$p < 0.05$, error bars represent ± SD

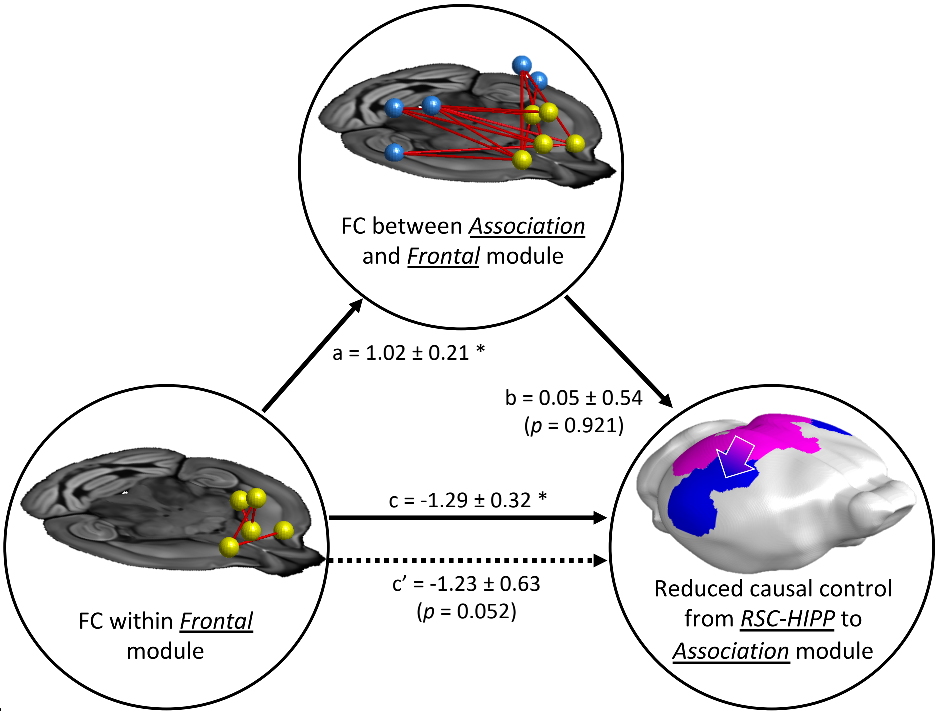

**Fig. S10** No mediation effect was observed from the dependent variable (enhanced FC within *Frontal* module) to the independent variable (decreased causal control from *RSC-HIPP* to *Association*) via mediator (enhanced FC between *Association* and *Frontal* modules).*$p < 0.001$.
